# Biallelic non-productive enhancer-promoter interaction precedes imprinted expression of *Kcnk9* during mouse neural commitment

**DOI:** 10.1101/2023.09.26.559498

**Authors:** Cecilia Rengifo Rojas, Jil Cercy, Sophie Perillous, Céline Gonthier-Guéret, Bertille Montibus, Stéphanie Maupetit-Méhouas, Astrid Espinadel, Marylou Dupré, Charles C. Hong, Kenichiro Hata, Kazuhiko Nakabayashi, Antonius Plagge, Tristan Bouschet, Philippe Arnaud, Isabelle Vaillant, Franck Court

**Affiliations:** Genetics, Reproduction and Development Institute (iGReD), CNRS, INSERM, Clermont Auvergne University, Clermont-Ferrand, France; Department of Medical and Molecular Genetics, King’s College, London, UK; Department of Medicine, University of Maryland School of Medicine, Baltimore; Department of Maternal-Fetal Biology, National Research Institute for Child Health and Development, 2-10-1 Okura, Setagaya, Tokyo 157-8535, Japan; Department of Human Molecular Genetics, Gunma University Graduate School of Medicine 3-39-22 Showa, Maebashi, Gunma, 371-8511 JAPAN; Department of Biochemistry, Cell and Systems Biology, Institute of Systems, Molecular and Integrative Biology, University of Liverpool, Liverpool, UK; Institut de Génomique Fonctionnelle, CNRS, INSERM, Université de Montpellier, Montpellier, France

**Keywords:** Genomic imprinting, chromatin looping, brain-specific expression, remote transcriptional control, Birk-Barel

## Abstract

How constitutive allelic methylation at imprinting control regions (ICRs) interacts with other levels of regulation to drive timely parental allele-specific expression along large imprinted domains remains partially understood. To gain insight into the regulation of the *Peg13-Kcnk9* domain, an imprinted domain with important brain functions, during neural commitment, we performed an integrative analysis of the epigenetic, transcriptomic and cis-spatial organisation in an allele-specific manner in a mouse stem cell-based model of corticogenesis that recapitulates the control of imprinted gene expression during neurodevelopment. We evidence that despite an allelic higher-order chromatin structure associated with the paternally CTCF-bound *Peg13* ICR, the enhancer-*Kcnk9* promoter contacts can occur on both alleles, although they are only productive on the maternal allele. This observation challenges the canonical model in which CTCF binding isolates the enhancer and its target gene on either side, and suggests a more nuanced role for allelic CTCF binding at some ICRs.

## Introduction

The functional specialisation of each cell and tissue type is of critical importance for higher multicellular organisms. It is based on the ability of the cell to respond to developmental and environmental cues by generating a specific gene expression profile. General principles governing this process are being identified. In particular, key regulatory DNA sequences, sequence-specific transcription factors, epigenetic modifications and the spatial organisation of the genome interact to regulate gene expression levels (*Shukla et al., 2022*), raising the question of how the coordinated action of these regulatory layers is orchestrated. For a particular set of genes in mammals, the imprinted genes, which are expressed in a parent-of-origin-specific manner, this issue is even more pressing.

Genomic imprinting is a key developmental process whereby some mammalian genes are expressed by only one allele, depending on their parental origin. Most of the 200 imprinted genes identified to date are involved in key biological processes such as cell proliferation, fetal and placental growth, energy homeostasis and metabolic adaptation (*Tucci et al, 2019*). Genomic imprinting also plays a central role in brain function and behaviour, with many imprinted genes being expressed only in neural lineages (*Ivanova and Kelsey, 2011*). Consequently, misregulation of imprinted genes is causally implicated in severe neurobehavioural disorders such as Prader-Willi and Angelman syndromes.

In both humans and mice, most imprinted genes are organised into evolutionarily conserved genomic clusters that contain between two and a dozen maternally and paternally expressed genes in regions spanning up to several megabases. Allele-specific expression in each of these clusters is primarily regulated by DNA methylation at discrete cis-acting regulatory elements known as imprinting control regions (ICRs). Each ICR overlaps with a differentially methylated region (DMR), which harbours allelic DNA methylation inherited from the male or female gamete and subsequently maintained throughout development (*i.e*: germline DMR). The resulting constitutive allelic DNA methylation at ICRs is critical for orchestrating allele-specific expression along the imprinted domain by influencing a combination of regulatory mechanisms, some of which are tissue-specific, leading to the complex and specific spatio-temporal expression pattern of imprinted genes (*Tucci et al, 2019*).

Specifically, histone modifications, cis-spatial organisation and tissue-specific regulatory regions have all been documented to contribute, along with DNA methylation, to this long-range ICR-mediated tissue-specific regulation of imprinted domains (*Noordemer and Feil*, 2020; *Hanna and Kelsey, 2021;*). Several studies at imprinted genes have correlated tissue-specific imprinted expression at imprinted genes with tissue-specific differences in histone modifications at their promoter region (*Yang et al., 2003; Lau et al., 2004; Li et al., 2004; Monk et al.,2006).* For example, placenta-specific paternal deposition of the repressive marks H3K9me2 and H3K27me3 contributes to placenta-specific maternal expression at a subset of genes in the mouse *Kcnq1* domain (*Umlauf et al., 2004, Wagschal et al., 2008; Mager et al., 2003*). In addition, at the maternally methylated ICRs, all of which are also promoters, timely developmental loss or gain of the repressive H3K27me3 mark on the paternal allele contributes to appropriate tissue-specific paternal expression (*Maupetit-Mehouas et al., 2016*). Allelic methylation at ICRs may also influence long-range chromatin interactions between enhancer and promoters along imprinted domain. At a genome-wide level, such interactions between regulatory elements and their target genes are facilitated by sub-chromosomal structures, the Topological Associated Domains (TAD) (*Dixon, 2012*). A study performed on *H19-Igf2* and *Dlk1-Gtl2*, two domains controlled by a paternally methylated ICR, showed that binding of the methyl-sensitive and boundary protein CTCF to the unmethylated allele of the ICR induces an allele-specific sub-TAD organisation, which is proposed to facilitate the establishment and maintenance of the imprinted transcriptional programme (*lleres, 2019*). This observation echoed previous studies showing that allelic methylation and allelic CTCF binding at *H19* ICR are both critical for mediating parent-specific chromatin loops and ensuring timely and allele-specific enhancer-promoter interaction along the *Igf2-H19* domain (*Murrell et al, 2004; Kurukuti, 2006; Court et al, 2011*). A recent study showing that allelic CTCF binding at a post-implantation DMR (secondary DMR) structures the *Grb10-Ddc* locus to direct the proper enhancer-promoter interactions in the developing heart further illustrates the interplay between methylation and CTCF binding to control instructive allelic chromatin configurations at imprinted loci (*Juan, 2022*). The observation that CTCF binds the ICR of multiple imprinted loci (*Lin et al., 2011*) suggests that allelic chromatin structure may be a commonly used strategy whereby ICRs direct mono-allelic expression along large genomic imprinted domains. For most imprinted clusters, however, this claim has yet to be formally evaluated.

Taken together, these observations highlight that deciphering how constitutive allelic methylation at ICRs can direct tissue- and stage-specific allele-specific expression along imprinted domains requires simultaneous analysis of the dynamics of multiple layers of regulation during cell identity acquisition. To gain insight into the regulation of the *Peg13-Kcnk9* domain during neural commitment, we precisely monitor the epigenetic, transcriptomic and cis-spatial organisation in an allele-specific manner in a mouse stem cell-based model of corticogenesis, which we have previously shown to recapitulate the in vivo epigenetic control of imprinted gene expression (*Bouschet et al., 2017*).

The *Peg13-Kcnk9* domain is an evolutionarily conserved domain with important functions in the brain. It contains 5 genes, two of which show conserved imprinting between humans and mice: the paternally expressed non-coding RNA *Peg13* and the potassium channel gene *Kcnk9*, which is maternally expressed specifically in the brain. The other three genes, *Trappc9*, *Chrac1* and *Ago2*, do not show imprinting in humans, whereas they are preferentially expressed by the maternal allele specifically in the mouse brain (*Smith et al., 2003; Ruf et al., 2007, Court, 2014; Babak et al., 2015; Bonthuis et al., 2015; Bouschet et al., 2016; Andergassen et al., 2017*). Mutations in these genes are associated with neurodevelopmental and neurological disorders (*Liang et al., 2020; Wilton et al., 2020; Aslanger et al., 2022; Lessel et al., 2020*), including Birk-Barel intellectual disability syndrome, which is caused by a maternally inherited mutation in *Kcnk9* (*Linden et al., 2007; Barel et al., 2008; Cooper et al., 2020*).

Little is known about the mechanisms that control expression along this domain. The *Peg13* promoter overlaps with a germline DMR and is proposed to be the ICR for the locus (*Smith et al., 2003; Ruf et al., 2007, Court, 2014)*. In an attempt to unravel the underlying mechanism, a study in human brain tissue supports that this DMR controls *Kcnk9* and *Peg13* imprinted expression through a CTCF-mediated enhancer-blocking activity (*Court et al., 2014*). However, a formal demonstration of this model and whether it accounts for the temporal imprinting of expression along the domain during neural identity acquisition remains to be established. Here, we have performed an integrative analysis of multiple levels of regulation, providing us with a comprehensive view of the molecular events that take place during the establishment of maternal *Kcnk9* expression during neural commitment. Our main observation challenges the canonical model of CTCF-mediated enhancer-blocking activity and suggests a more nuanced role for allelic CTCF binding at the ICR of this locus.

## Results

### *Kcnk9* gains maternal expression upon neural commitment

Using a microfluidic quantitative RT-PCR approach, we observed, in agreement with allelome studies (*Babak et al., 2015; Andergassen et al., 2017*), that *Peg13* is expressed in a wide range of adult mouse tissues and at different stages of development, with a predominant expression in the brain. In contrast, *Kcnk9* expression is restricted to brain tissues (Supplementary Figure 1A). Expression analyses conducted on brain isolated from reciprocal crosses of C57BL/6J and Mus musculus mollosinus (JF1) newborn F1 hybrids, further confirmed that *Peg13* is paternally expressed and *Kcnk9* maternally expressed (Fig 1A). To investigate the mechanisms responsible for this brain-specific imprinted expression, we adapted a stem cell-based model of corticogenesis, which we have previously shown to recapitulate the in vivo epigenetic control of imprinted gene expression (*Bouschet et al., 2017*), to mouse ESC lines we derived from reciprocal crosses between C57BL/6 (B6) and JF1 mouse strains (hereafter; B/J and J/B). Reciprocal crosses allow us to interrogate the parental allelic origin using informative polymorphisms (SNPs).

**Figure 1:**
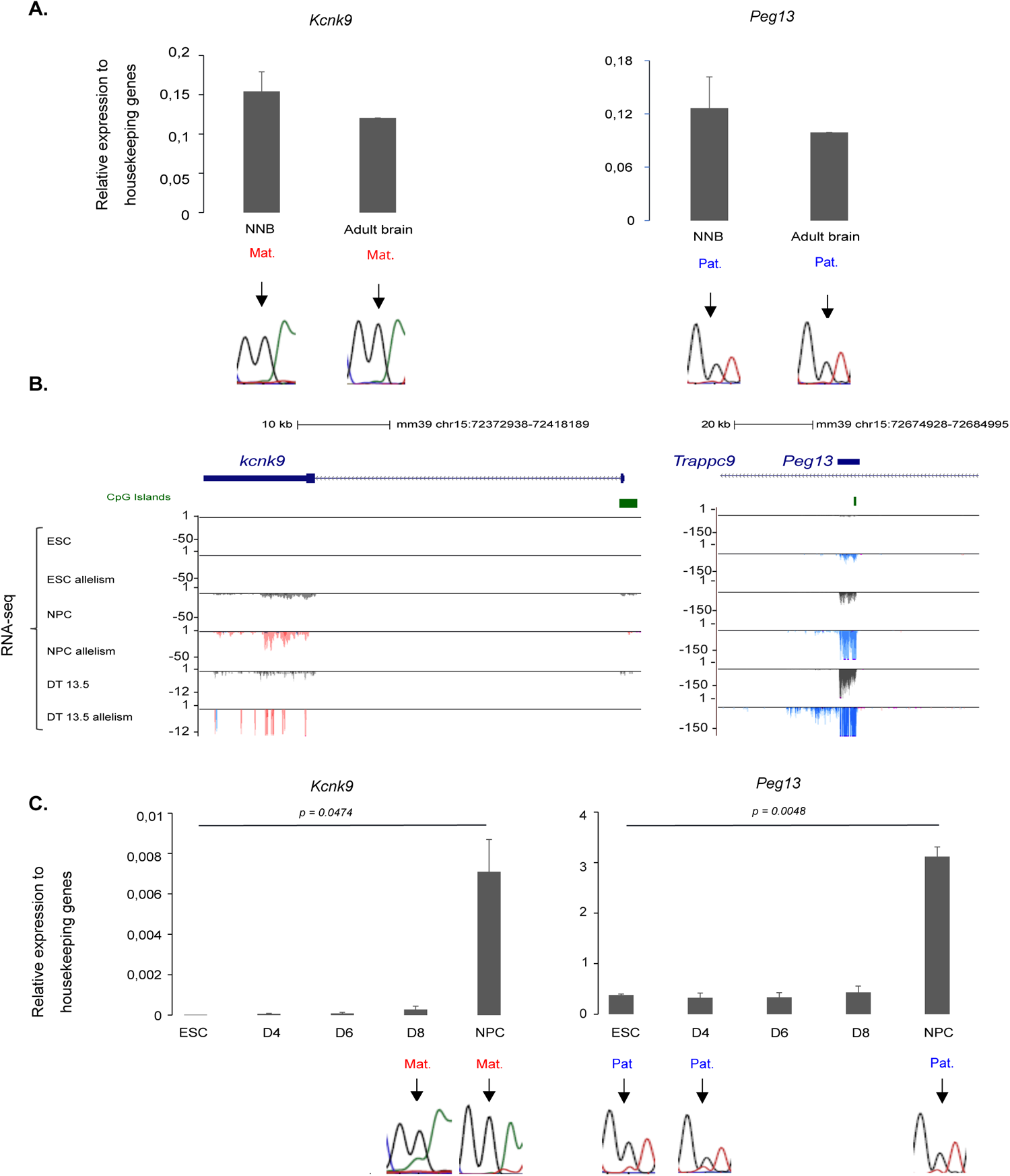
*Kcnk9* and *Peg13* expression dynamics during neural commitment. **A)** Quantitative RT-PCR analysis to assess the expression levels of *Kcnk9* and *Peg13* in the brain of newborn (NBB; n=4) and adult (n=2) mice. The parental origin of expression is shown in the lower panel. **B)** Genome browser view at the *Kcnk9* and *Peg13* loci to show the allelic oriented RNA-seq signal in ESC, NPC and embryonic brain (Dorsal Telencephalon ((DT); re-analysed data from Bouschet et al., 2016). For each condition, the quantitative and merged parental allelic RNA-seq signal is shown in the top and bottom panels, respectively. Maternal and paternal expression are shown in red and blue, respectively. **C)** Quantitative RT-PCR analyses to assess *Kcnk9* and *Peg13* expression levels in ES cells (n = 4) and at day 4 (D4; n = 2), D6 (n=2), D8 (n = 2) and NPC (D12; n=4) stages of in vitro corticogenesis. The parental origin of expression is shown in the lower panel. Statistical significance was determined with the unpaired t test (p values in the figure). In **A)** and **C)** the results are presented as percentage of expression relative to the geometric mean of the expression of the three housekeeping genes *Gapdh*, *Gus* and *Tbp*. The data are presented as the mean ± SEM. The parental origin of expression was determined by direct sequencing of sample-specific PCR products incorporating a strain-specific SNP in the regions analysed, representative data are shown.

We focused on the first 12 days of in vitro corticogenesis. During this period, ES cells predominantly differentiated into Neural precursor cells (NPC), as indicated by the marked down-regulation of the pluripotency marker *Pou5f1* and the up-regulation of the neural precursor markers *Nestin* and *Pax6* (Supplementary Figure 1B). RNA-seq and RT-qPCR approaches showed that in ES cells, imprinted expression is restricted to the *Peg13* gene, which shows weak paternal expression. Upon differentiation towards NPC, paternal *Peg13* expression is upregulated while *Kcnk9* gains maternal expression (Figure 1B,C). During this time window, the other three genes of the domain, *Trapc9*, *Ago2* and *Chrac1*, are biallically expressed (Supplementary Figure 1C). This pattern is reminiscent of that observed in primary NSCs (neurospheres) cultured from newborn mice (*Claxton et al., 2022*). In addition, re-analysis of RNA-seq from the E13.5 dorsal telecephalon (*Bouschet et al., 2016*) further demonstrated that the imprinted expression pattern obtained at the *Peg13* domain in NPC from our corticogenesis model system, restricted to *Kcnk9* and *Peg13*, recapitulates those observed in the embryonic brain in vivo (Fig 1B, Supplementary Figure 1D).

Our stem cell-based model of corticogenesis to generate NPC thus provides a relevant framework to uncover mechanisms acting at the *Peg13* domain, and in particular those involved in the imprinted expression of *Peg13* and *Knck9*, in in vivo NSC and during early brain development.

### Peg13 DMR maternal methylation is required for *Kcnk9* maternal expression

Peg13 DMR is the putative ICR proposed to control the imprinted expression of the entire locus. To more formally evaluate the role of allelic methylation of this region in the regulation of *Kcnk9*, we assessed expression in the upper part, enriched in brain tissue, of *Dnmt3L*^-/+^ E9.5 embryos derived from *Dnmt3L*^-/-^ females, in which DNA methylation imprints at ICRs are not established during oogenesis (*Bourc’his et al., 2001; Hata et al., 2002*). In wild-type embryos, we confirmed paternal and maternal expression of *Peg13* and *Kcnk9*, respectively (Figure 2). In mutant embryos, the lack of maternal DNA methylation at the *Peg13* DMR (Supplementary Figure 2) results in increased and biallelic expression of *Peg13*, while *Kcnk9* expression is lost (Figure 2). This observation supports that the *Peg13* DMR is the ICR of the locus and indicates that its maternal methylation is required for the maternal expression of *Kcnk9*.

**Figure 2:**
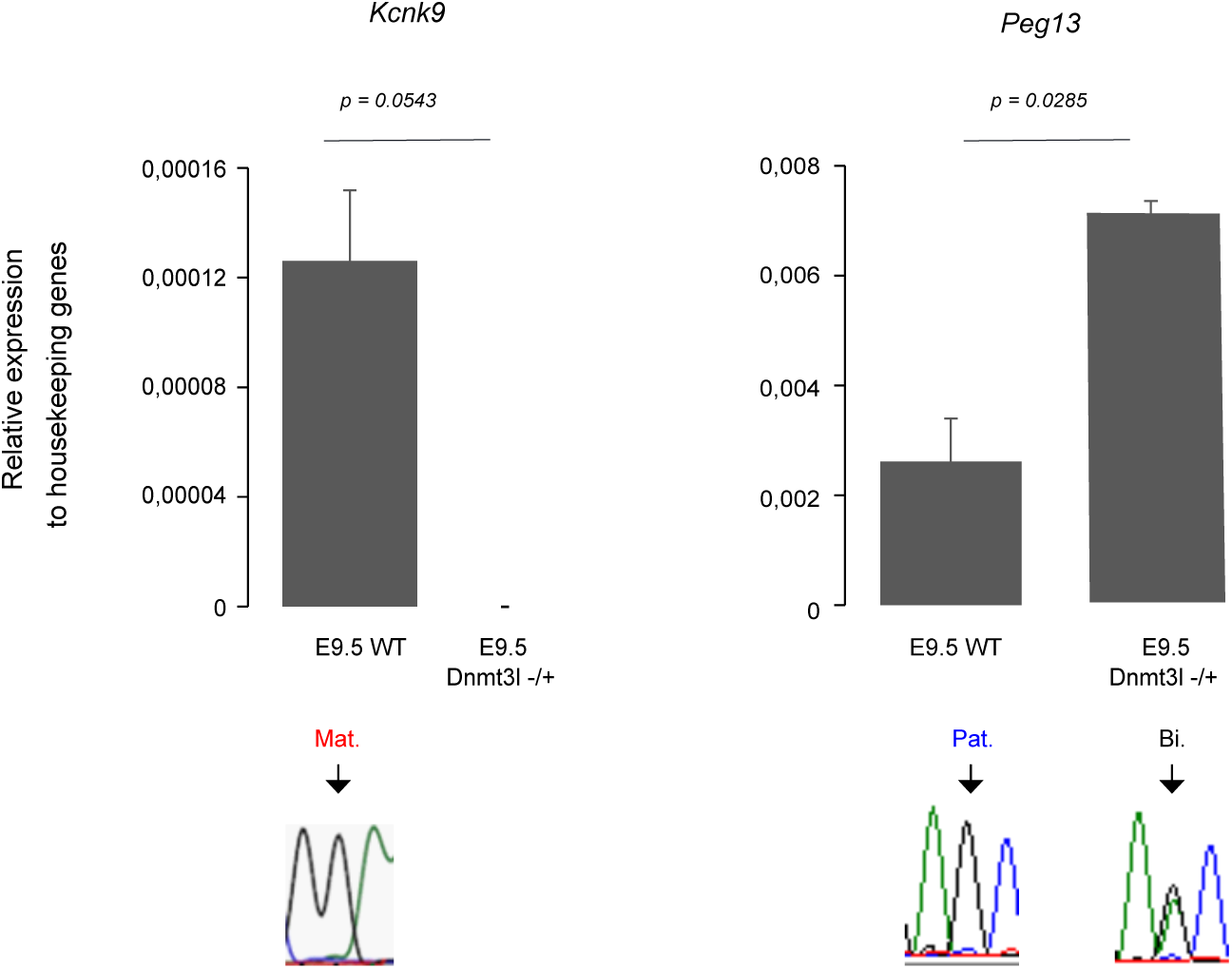
*Kcnk9* expression is lost in *Dnmt3l*−*/*+ E9.5 embryos. Quantitative RT-PCR analysis to assess *Kcnk9* and *Peg13* expression levels in the upper part of WT (n=2) and *Dnmt3l*−*/*+ (n=3) E9.5 embryos. Statistical significance was determined with the unpaired t test (p values in the figure). The data are presented as the mean ± SEM. The parental origin of expression is shown on the lower panel.

### Changes in imprinted expression upon neural commitment are not associated with changes in epigenetic signatures at *Peg13* DMR

We then performed an integrative analysis based on allelic ChIP-seq, Cut & Run (C&R) and ChIP-qPCR approaches coupled with non-allelic data mining derived ATAC-seq and WGBS, to evaluate whether changes in the epigenetic signature at the *Knck9* and *Peg13* promoters can account for their change in expression upon differentiation towards NPC.

In ES cells, the *Peg13* promoter showed the characteristic feature of an ICR (*Henckel & Arnaud, 2010*) with DNA methylation, the repressive histone mark H3K9me3 and the zinc finger protein ZFP57 associated with its maternal allele, while the permissive histone marks H3K4me2, H3K4me3 and H3K27ac are associated with its paternal allele (Fig 3, Supp fig 3). This allelic signature is maintained in the NPC, where the permissive histone marks are more widely distributed along the gene on the paternal allele, and further in the neonatal mouse brain with no major change in level despite *Peg13* up-regulation (Figure 1, Figure 3, Supplementary Figure 3).

**Figure 3:**
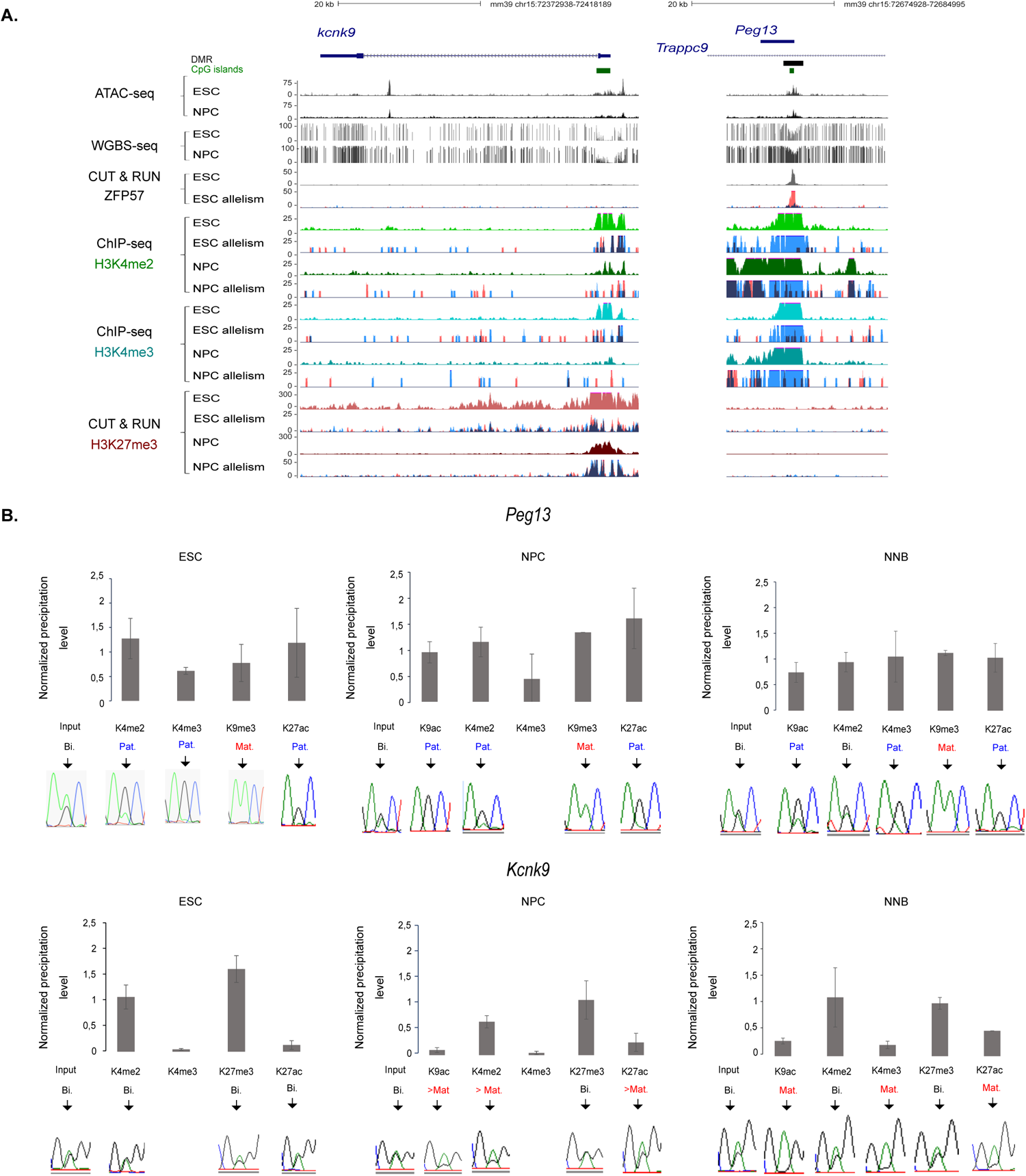
Epigenetic signatures at *Kcnk9* and *Peg13* in ESC, NPC and neonatal brain. **A)** Genome Browser view at the *Kcnk9* and *Peg13* loci to show CpG island (CGI) positions and ATAC-seq, methylation (WGBS), ZPF57, H3K4me2, H3K4me3 and H3K27me3 enrichments in ESC and NPC. For ZFP57 and the histone marks, the quantitative and the merged parental allelic signal are shown in the upper and lower panels, respectively. Maternal and paternal enrichment are shown in red and blue respectively. **B)** Chromatin analysis following native ChIP-qPCR to analyse the deposition of the indicated histone marks at *Peg13* and *Kcnk9* promoters. The precipitation level was normalized to that obtained at the *Rpl30* promoter (for H327ac, H3K4me2 and H3K4me3), the *HoxA3* promoter (for H3K27me3) and IAP (for H3K9me3). For each condition, values are the mean of independent ChIP experiments (*n*), each performed in duplicate: ESC (*n* =3); NPC (*n* = 3); neonatal brain (*n* = 5). The data are presented as the mean ± SEM. The allelic distribution of each histone mark was determined by direct sequencing of the sample-specific PCR products containing a strain-specific SNP in the analysed region, representative data are shown.

The *Kcnk9* promoter is localised to a CpG island that remains unmethylated in both ES cells and NPC, and further in embryonic and adult brain tissues, highlighting that *Kcnk9* is not controlled by methylation dynamic in its promoter (Fig 3A, Supp figure 3B). In contrast to *Peg13*, no allelic signature is observed at the *Kcnk9* promoter in ES cells. A broad domain of H3K27me3 biallelically marks the gene body. This repressive mark is further associated with the permissive H3K4me2 on both alleles of the promoter, forming a bivalent signature proposed for poise gene expression (*Kumar et al., 2021*). In NPC, H3K27me3 is lost from the gene body, while the promoter retains bivalency, albeit with lower levels of H3K27me3 (Figure 3A,B). ChIP-qPCR also showed that the slight increase in H3K9ac and H3K27ac, marks associated with active transcription, occurs preferentially on the maternal allele (Figure 3B). This trend is further enhanced in the neonatal brain, where H3K9ac, H3K27ac and H3K4me3 mark the maternal allele of the promoter, while the biallelic bivalent signature H3K4me2/H3K27me3 is maintained (Figure 3B).

These observations support that the upregulation of paternal *Peg13* expression and the gain of maternal *Kcnk9* expression in neural commitment are not driven by changes in the *Peg13* DMR/putative ICR epigenetic signature. Specifically, for *Kcnk9*, maternal expression is induced despite the presence of H3K27me3 in the promoter and is accompanied by biallelic loss of H3K27me3 in the gene body and a gain of permissive/activating, mainly acetylation, marks on the maternal promoter reflecting transcriptional activity.

### Biallelic interactions between the *Kcnk9* promoter and its putative regulatory region in ES cells precede the maternally biased interaction in NPCs

In addition to epigenetic modifications, higher-order chromatin structure through chromatin looping is another layer of regulation that controls expression along imprinted clusters and facilitates enhancer-promoter interaction within topologically associated domain (TAD). The DNA-binding protein CTCF is a key determinant in the formation of these loops and is frequently found at their base.

Allelic C&R analyses showed that in both ES and NPC, the *Peg13* DMR/putative ICR is tightly bound by CTCF in a paternal-specific manner, whereas CTCF binds both alleles of the *Kcnk9* promoter and the 3’ edge of its unique intron (Figure 4).

**Figure 4:**
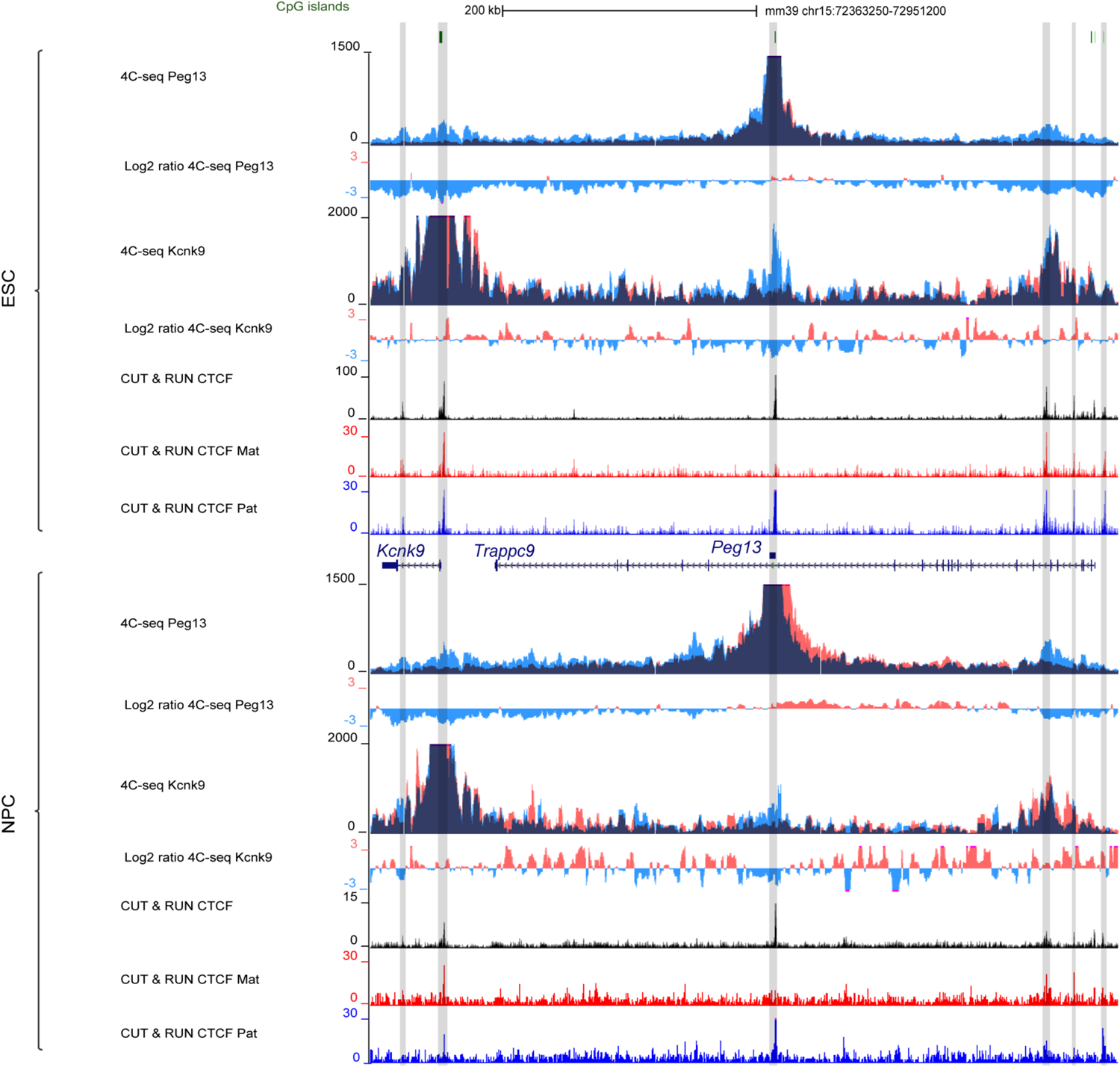
*Peg13* DMR and *Kcnk9* promoters interact with the same putative enhancer in ESC and NPC. Genome Browser view on the *Kcnk9* -*Trappc9* genomic region to show in ESC (upper panel) and NPC (lower panel) allelic 4C-seq, for the Peg13 DMR and Kcnk9 promoter viewpoints, and CTCF C&R signals. 4C-seq are shown by merging the allelic signals, contacts mediated by the paternal and maternal alleles are shown in blue and red, respectively. The ratio of maternal/paternal interactions is indicated. CTCF-bound regions are highlighted. Binding is biallelic for all except the paternally bound *Peg13* DMR.

To assess whether these regions form chromatin loops and to identify putative distant regulatory regions, we performed allelic 4C-seq using the *Peg13* DMR and the *Kcnk9* promoter as viewpoints. While the *Peg13* domain is all contained within a larger TAD (as defined in cortex by *Dixon et al., 2012*), the signal obtained for *Peg13* DMR in ES cells was largely restricted to the imprinted domain, from downstream *Kcnk9* to upstream *Ago2* (Supplementary Figure 4). Strikingly, and in the two reciprocal crosses, it was exclusively mediated by the paternal unmethylated *Peg13* DMR, highlighting that paternal CTCF binding promotes higher-order chromatin structure differences between the two parental alleles (Figure 4, Supplementary Figure 4 and see next section). Although paternal-specific contacts were observed along the entire imprinted domain, stronger signals were detected at *Kcnk9*, centred around the CTCF-bound promoter and 3’ edge of the intron, and at a biallelic CTCF-bound region in the 5’ part of *Trappc9* (Figure 4, Supplementary Figure 4). This second signal peaks in a *Trappc9* intron previously identified as a putative regulatory region controlling tissue-specific expression of the domain (*Claxton et al., 2022*). These paternal-specific contacts are mainly maintained in NPC, where the interaction with the *Trappc9* intronic putative regulatory region is reinforced.

The same analysis using the *Kcnk9* promoter as view point confirmed that it interacts with *Peg13*DMR only on the paternal allele in ES cells and, although weaker, in NPC. In addition, the *Kcnk9* promoter also interacts with the intronic putative regulatory region in *Trappc9*, from both alleles in ES cells and preferentially from the maternal allele in NPC (Figure 4).

These results identified a putative regulatory region, hereafter referred to as PE (Putative Enhancer), that interacts with the paternal *Peg13* promoter in ES and NPC, and preferentially with the maternal *Kcnk9* promoter in NPC, and is thus a candidate for regulating the imprinted expression of both genes during neural commitment. However, contrary to expectation, contacts with the *Kcnk9* promoter are already established in ES cells and from both alleles. This supports that non-productive biallelic contacts between the *Kcnk9* promoter and the putative regulatory region in ES cells precede maternally biased productive contacts in NPC.

### Contacts from PE and Peg13DMR structure the higher-order chromatin conformation in the *Peg13* domain

To further investigate the extent to which the intronic PE influences conformation along the *Peg13* domain, we performed allelic 4C-seq using this region as a viewpoint and visualised the data together with allelic 4C-seq data for *Peg13* DMR and *Kcnk9* promoter, respectively and re-analysed high-resolution but non-allelic Hi-C data in ESCs and NPC in vivo (*Bonev et al., 2017*).

Hi-C data revealed that the *Peg13* imprinted domain resides in two sub-TADs that are conserved in ESC and NPC (Figure 5). The centromeric sub-TAD is anchored to CTCF-bound regions in the 5’ part of *Trappc9* and in *Kcnk9,* presumably in the intron (the 5 kb resolution of the Hi-C data do not allow to precisely map the boundaries regions), respectively, thus isolating *Kcnk9* and *Peg13* from *Chrac1* and *Ago2* localised in the telomeric sub-TAD. *Trappc9* promoter being localized at the boundary between these two sub TADs (Figure 5, Supplementary Figure 5). Furthermore, in line with 4C-seq data, paternal CTCF binding at the *Peg13* DMR subdivided the telomeric sub-TAD into two sub-domains, presumably on the paternal allele only, in a structure maintained in ES cells and NPC *(Fig. 5*).

**Figure 5:**
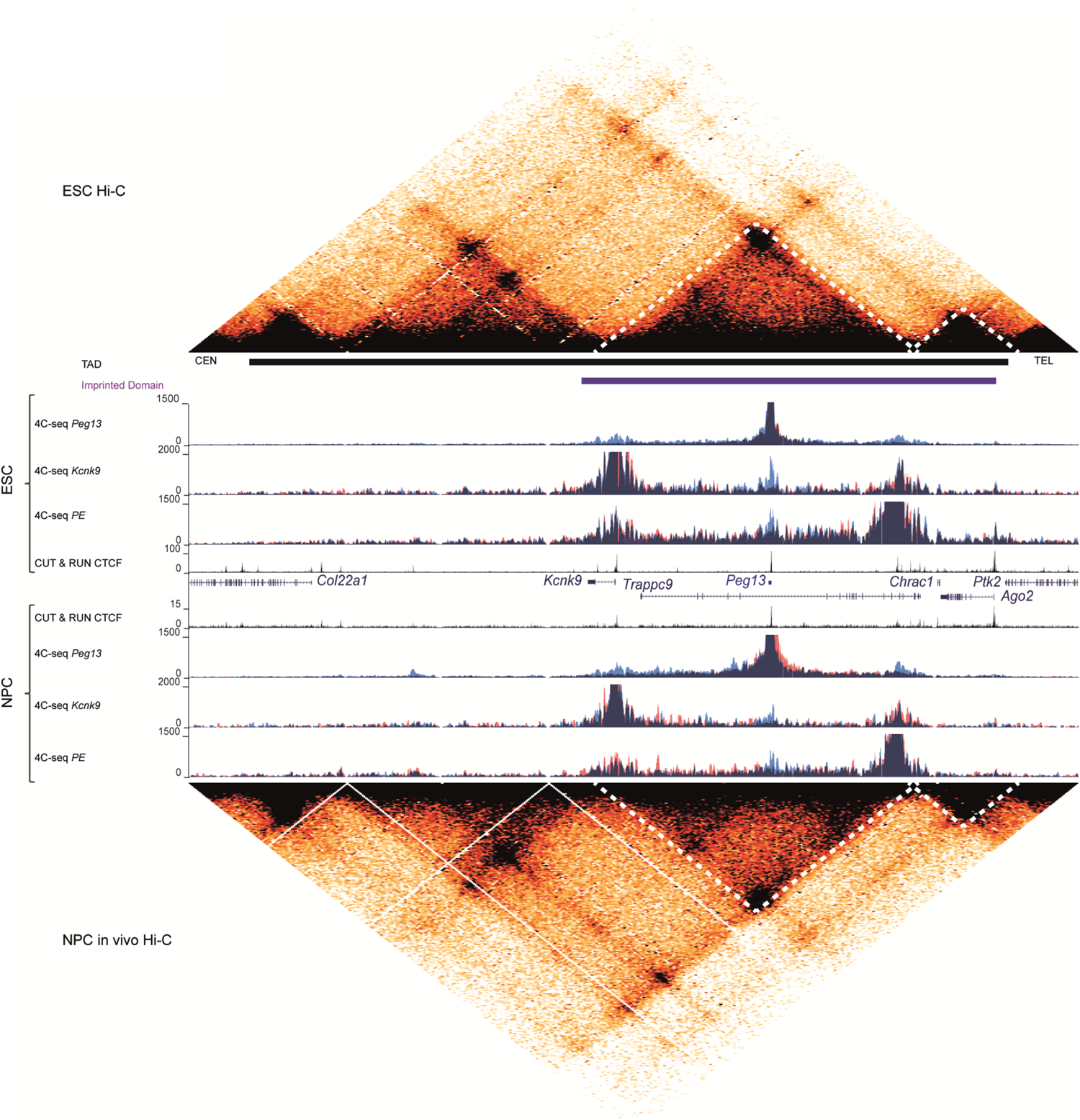
*Peg13* DMR and *PE* interactomes structure the higher-order chromatin conformation at the *Peg13* domain. Genome browser view at the *Peg13* domain containing TAD to show in ESC (top panel) and NPC (bottom panel); re-analysed HiC data, allelic 4C-seq for the *Peg13* DMR, *Kcnk9* promoter and the PE viewpoints and and CTCF C&R signals. 4C-seq are shown by merging the allelic signals, contacts mediated by the paternal and maternal alleles are shown in blue and red, respectively. Sub-TADs dividing the imprinted domain are indicated by a dotted line.

Interestingly, in ES cells, the ability of this higher-order chromatin structure to isolate Sub-TAD or domains from each other is not fully functional. The 4C-seq signal obtained for the PE region was indeed mainly, but not entirely, restricted to the centromeric sub-TAD. It was also observed, albeit to a lesser extent, in the telomeric region, including at the *Ago2* promoter (Figure 5). In addition, PE strongly contacts the *Kcnk9* promoter from both alleles, despite the *Peg13*-DMR associated subdomains on the paternal allele (Figure 5, Supplementary Figure 5).

In NPC, PE contacts were restricted to the centromeric sub-TAD, which coincides with a stronger interaction at the sub-TAD boundary (arrow b in sup Fig. 5), which may enhance its insulating capacity (Figure 5, Supplementary Figure 5). Along this centromeric sub-TAD the pattern observed in ES cells remains largely stable in NPC, with a conserved, albeit weaker, contact from the paternal allele with the *Peg13* DMR. In addition, the nature of the strong contact with the *Kcnk9* promoter, also observed in the Hi-C data (arrow *a* in supp figure 5), changes from biallelic in ES cells to preferentially from the maternal allele in NPC. This change occurs along the entire sub-TAD, with preferential maternal and paternal interactions, respectively, with the regions located on either side of the *Peg13* DMR (Supplementary Figure 5).

These data mirror and support those obtained from the 4C-seq analysis using the *Peg13* DMR and *Kcnk9* promoter as viewpoints (Figure 4& 5). They highlighted that the *Peg13* DMR structures the centromeric sub-TAD into two paternal sub-domains, isolating the *Kcnk9* promoter from the putative enhancer (PE) on the paternal allele. However, this structure is likely to be circumvented in ES cells where PE strongly contacts *Kcnk9* from both alleles, further indicating that these contacts precedes *Kcnk9* imprinted expression. In NPC, and consistent with the gain of maternal expression at *Kcnk9*, contacts between PE and the promoter now occurs preferentially, although not exclusively, on the maternal allele (Figure 5, Supplementary Figure 5).

### PE shows features of a biallelically active enhancer from ES to NPC and in the neonatal brain

The PE intronic region is one of the enhancers annotated by mouse transgenic experiments (*Visel et al., 2010*) with activity in mouse brains (dataset ID: mm1679 in Vista Enhancer Browser). This observation, combined with our chromatin structure studies, supports that this intronic region could be an enhancer for both *Peg13* and *Kcnk9*, but that the contacts already established in ES cells are not sufficient per se to induce maternal *Kcnk9* expression and paternal upregulation of *Peg13*.

We therefore performed an integrative analysis to assess whether the changes in the molecular signature at this region could account for the change in expression of *Kcnk9* and *Peg13* between ES and NPC. Non-allelic ATAC-seq and WGBS datasets indicated that this region is in an open chromatin configuration, with a strong ATAC-seq signal and depletion for DNA methylation in both cell types and brain tissue (Fig. 6A, sup Fig 6). Allelic C&R and ChIP-qPCR further demonstrated that in both ES cells and NPC, as well as in neonatal brains, several permissive/activating marks, including H3K27ac, a signature of active enhancers, are enriched on both alleles, whereas the repressive H3K27me3 is absent. In addition to the histone signature, active enhancers have also been shown to produce non-coding RNA called eRNA (*Kim et al., 2010*). Refined analysis of allelic RNA-seq precisely identified a biallelically expressed RNA originating from this region in NPC and embryonic brains (Fig. 6B, left part). Time course analysis by RT-qPCR confirmed that this RNA is biallelically expressed and that its expression, although low, gradually increases as ES cells differentiate into NPC and is maintained in neonatal brains (Fig. 6B).

**Figure 6:**
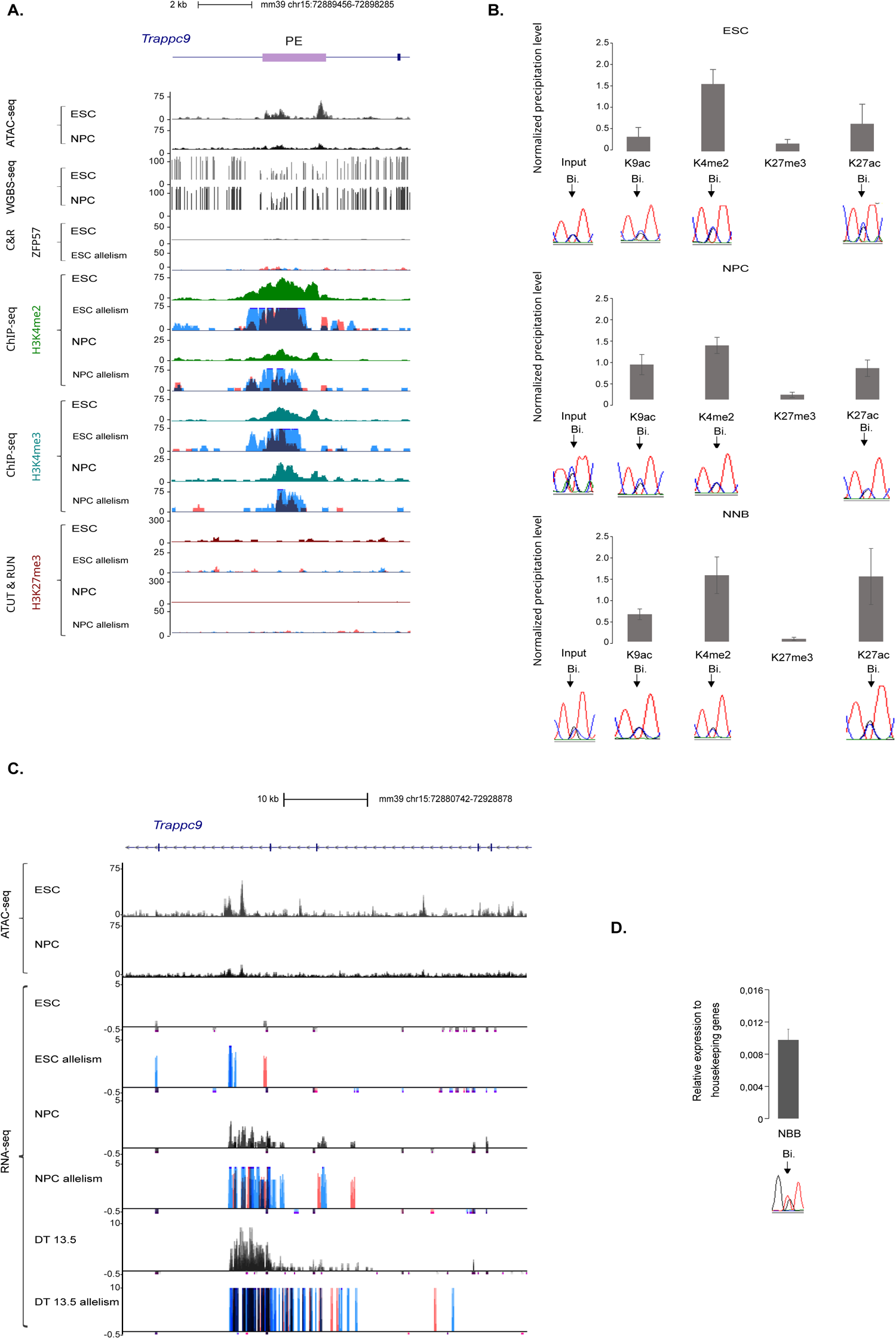
Dynamics of PE molecular signature in neural commitment. **A)** Genome browser view at the *intronic PE region* to show CpG island (CGI) position and ATAC-seq, methylation (WGBS), ZPF57, H3K4me2, H3K4me3 and H3K27me3 enrichments in ESC and NPC. For ZFP57 and the histone marks, the quantitative and the merged parental allelic signal are shown in the upper and lower panels, respectively. Maternal and paternal enrichment are shown in red and blue respectively. **B)** Chromatin analysis following native ChIP-qPCR to analyse the deposition of the indicated histone marks. The precipitation level was normalised to that obtained at the *Rpl30* promoter (for H3K27ac, H3K4me2 and H3K4me3), the *HoxA3* promoter (for H3K27me3) and IAP (for H3K9me3). For each condition, values are the mean of at least three independent ChIP experiments (*n*), each performed in duplicate: ESC (*n* = 4); NPC (*n* = 3); neonate brain (*n* = 4). The allelic distribution of each histone mark was determined by direct sequencing of the sample-specific PCR products containing a strain-specific SNP in the analysed region, representative data are shown. **C)** Genome browser view at the *PE region* to show the allelic oriented RNA-seq signal in ESC, NPC and embryonic brain (Dorsal Telencephalon ((DT); reanalysed data from Bouschet et al., 2016). For each condition the quantitative and the merged parental allelic RNA-seq signal is shown in the top and bottom panels, respectively. Maternal and paternal expression are shown in red and blue respectively. **D)** Quantitative RT-PCR analyses to assess PE-associated eRNA expression in neonatal brains (n=4). The parental origin of expression is shown on the lower panel. **B)** and **D)** The data are presented as the mean ± SEM.

Taken together, these observations support that the CTCF-bound region in the *Trappc9* intron is a bona fide biallelic enhancer that is pre-loaded in an active, but not productive, configuration on the *Kcnk9* promoter in ES cells. Increased activity during NPC formation and further in the neonatal brain is associated with an increase in biallelic e-RNA production.

Notably, we also observed that e-RNA expression is maternally biased in the adult brain (Supplementary Figure 6) and, by data mining, that H3K27ac is enriched on the maternal allele of the PE region in the adult mouse frontal cortex (Supplementary Figure 6). This observation suggests that the activity of the enhancer can switch from a biallelic to a maternal bias in the adult brain.

## Discussion

In this study, we aimed to gain insight into the regulation of the *Peg13-Kcnk9* domain during neural commitment. Our mouse stem cell-based model of corticogenesis combined with integrative analyses of multiple layers of regulation provide us with a comprehensive view of the molecular events that take place along with the establishment of *Kcnk9* maternal expression. We have evidence that despite an allelic higher-order chromatin structure associated with CTCF, enhancer-*Kcnk9* promoter contacts can occur on both alleles, although they are only productive on the maternal allele. This observation challenges the canonical model in which CTCF binding acts as a chromatin boundary and suggests a more refined role for allelic CTCF binding at the gDMR and the resulting allelic chromatin loops at this locus.

The molecular patterns detected at the *Peg13* locus using our stem cell-based model of corticogenesis mimic the in vivo observations. This consistency between these results for imprinted expression, the chromatin signature or conformation and also for eRNA production is further evidence, that the in vitro corticogenesis used here is a powerful model that recapitulates the complex regulations that occur in vivo during early brain development (*Bouschet 2017*). Furthermore, our observations are consistent with those identified in a study conducted in human brain tissue (*Court et al., 2014),* suggesting that the underlying mechanisms of the regulation of the *Peg13-Kcnk9* domain are evolutionarily conserved. Our study thus provides a frame to further decipher the etiology of neurodevelopmental and neurological disorders associated to this locus. In particular, in addition to the documented missense mutation in *Kcnk9* (Linden et al., 2007; Barel et al., 2008; Cooper et al., 2020), the enhancer appears to be another key region to screen for mutations and/or epigenetic alteration in patients with suspected Birk-Barel.

The observation that *Kcnk9* and *Peg13* DMR are located in a different sub-TAD from *Trappc9*, *Chrac1* and *Ago2* in ES and NPC provides a framework to explain the absence of imprinting for these three genes at these stages and more globally in the developing brain. Further analysis is required to determine if and how this higher-order chromatin structure is reorganised during later brain development to allow the *Peg13* DMR to direct the mechanism by which these three genes switch from biallelic to preferential maternal expression in the postnatal brain. However, the recent proposition that enhancer-promoter interaction may be memorized to influence promoter activity later in development (*Xiao et al., 2021; Wurmser & Basu, 2022*) questioned whether the inter-SUB-TAD interactions observed between the enhancer and *Ago2* promoter in ES cell may not have a functional role to instruct later imprinted expression.

Consistent with its germline DMR status (*Ruf et al., 2007*) our data support that the *Peg13* promoter is the ICR of the locus. We have evidence that it exhibits the characteristic allelic molecular feature of an ICR and that its maternal methylation is required to control the maternal expression of *Kcnk9*, located approximately 250kb away. In addition, it is unique region of the domain that recruits CTCF in a parental allele-specific manner. It is reasonable to assume that the resulting differences in higher-order chromatin structure between the two parental alleles provide a framework within which this ICR can impose the imprinted transcriptional programme along the domain, as observed at *H19-Igf2* and *Dlk1-Gtl2* imprinted loci (*Lleres, 2019*).

However, the underlying mechanism lies outside the canonical model framework which postulates that parent-specific chromatin loops mediated by CTCF restrict enhancer-promoter interaction to the expressing allele only (*Murrell et al., 2004; Kurukuti, 2006; Lleres, 2019*). As previously documented for a minority subset of enhancer-promoter pairs (*Bonev et al. 2017*; *Gavi-Helm, 2014*), *Kcnk9* promoter-enhancer interactions are pre-established in non-expressing ES cells. The observation that there is no *Kcnk9* expression at this stage, despite the enhancer active signature, is intriguing. This may be explained by a cell context-dependent dual function for this regulatory element. As observed for other human and mouse regulatory elements (*Huang and Ovcharenko, 2022; I Simeoni et al., 2021*), it can recruit repressor or activator factors in ES cells and neural cells, respectively. More surprisingly, the enhancer -promoter contacts occur from both alleles, although they are only productive from the maternal allele after differentiation.

These observations suggest an interplay between the pre-existing chromatin structure, the allelic CTCF binding at *Peg13* DMR and the transcriptional machinery to shape imprinted expression during differentiation. In this model, CTCF-anchored loops provide a structural, but as yet uninformative, higher-order chromatin structure in ES cells. Upon differentiation and recruitment of ad hoc activator transcription factors to the enhancer, this framework guides productive contacts. On the paternal allele, the pre-existing interactions between the *Peg13* DMR and both the enhancer and *Kcnk9* promoter allow the gain of *Peg13* expression and keep *Kcnk9* silent. Specifically, we propose that rather than isolating the enhancer from the promoter, the CTCF-mediated loops here induce a three-way *Kcnk9* promoter-*Peg13* promoter-enhancer contact, where promoter competition for transcription factors and/or a physical barrier formed by the *Peg13* DMR between the *Kcnk9* promoter and enhancer keep *Kcnk9* silent. Such a multi-way interaction between enhancers and promoters, facilitated by ordered chromatin structure, has been proposed to regulate temporal expression along the α-globin locus. (*Marieke-Oudellar et al., 2019; Topfer et al. 2022*). In addition, there are several examples of an active promoter between an enhancer and another promoter that reduces the activity of the distal promoter (*Bartman et al., 2016; De Gobbi et al., 2006; Cho et al., 2018*). The absence of any specific interaction on the paternal allele other than with the *Peg13* DMR and the absence of a strong repressive signature, such as methylation, on its promoter rules out the action of a silencer and suggests that this mechanism is the main one to maintain *Kcnk9* silencing. Notably, the allelic *Peg13* DMR-associated subdomains and enhancer-*Kcnk9* promoter interactions we detected on the paternal allele are explained by such a three-way contact. On the maternal allele, the sole pre-existing interaction between the enhancer and *Kcnk9* promoter induce its maternal expression. Importantly, the associated recruitment of RNAPolII, which can promote enhancer-promoter interaction (*Zhang et al., 2023*), will in turn affect chromatin structure by strengthening this interaction, which becomes stronger on the maternal allele as expression increases. This model, which remains to be validated, explains the allelic specificity of the enhancer despite biallelic interactions with *Kcnk9* and provides an alternative to the canonical isolation model.

Our observation supports that although CTCF binds many ICRs to their unmethylated allele (*Prickett et al., 2013*), its function may not be universal between imprinted loci, where it may act through different mechanisms.

## Acknowledgements

We thank all members of P.A.’s team for critical reading of the manuscript. This research has been financed by the French government IDEX-ISITE initiative 16-IDEX-0001 (CAP 20–25) (Projet Emergence, to PA and FC, and projet Challenge 3-recherche to PA).

## AUTHORS CONTRIBUTIONS

Conceptualization, P.A, I.V, F.C; Experimental design, P.A, F.C; Data production, C.R.R, J.C, S.P, C.G.G, B.M, S.M.M, A.E, M.D, K.H, K.N, A.P, T.B, P.A, I.V, F.C; Data analysis, C.R.R, J.C, S.P, C.G.G, B.M, S.M.M, P.A, I.V, F.C Resources, C.C.H, K.H, K.N; Writing – Original Draft P.A; Writing –Review & Editing P.A, I.V, F.C; Visualization C.R.R, J.C, F.C; Supervision P.A, F.C; Project Administration P.A; Funding Acquisition P.A, F.C

## MATERIALS AND METHODS

### Cell culture and ES cell differentiation

The hybrid ES cell lines, both male, were previously derived from blastocysts obtained from crosses between C57BL/J (B) and JF1 (J) mice (Montibus et al., 2021) and were maintained in gelatin-coated dishes with ESGRO complete plus medium (Millipore, SF001-500P) containing LIF, BMP4, and a GSK3-β inhibitor. In vitro corticogenesis was performed as previously described (*Gaspard et al., 2009*), except that ES cells were plated on Matrigel-coated dishes (human ES cell-qualified matrix, Corning), and that the Defined Default Medium was supplemented with B27 (without vitamin A, Gibco) to improve cell survival, and with 1 μM dorsomorphin homolog 1 (DMH1) (purified by C.C.H) to promote neurogenesis (*Nelly et al., 2012*). Using this protocol, neural precursors (NP) cells are the main cell population after 12 days (D12) of in vitro corticogenesis (*Gaspard et al., 2009*).

### Material collection

Neonatal and adult brains were obtained from reciprocal crosses of C57BL/6J (B6) to Mus musculus molossinus JF1/Ms mice, (B6xJF1) F1 and (JF1xB6) F1 mice, referred to as ‘BJ’ and ‘JB’ in the text. Dnmt3L^-/+^ mouse embryos were generated by crossing homozygous Dnmt3L^-/-^ females (129SvJae-C57BL/6 hybrid genetic background) to WT JF1 male mice (M. musculus molossinus). E9.5 Dnmt3L^-/+^ conceptuses were collected from pregnant dams. Tail DNA was used for genotype analysis by PCR as previously described (*Hata et al., 2002*).

### DNA methylation Analysis

#### DNA extraction and bisulfite sequencing

DNA extraction was done as previously described (*Arnaud et al., 2006*). Bisulfite conversion was performed by using the EZ DNA methylationTM Gold Kit from Zymo (ref. D5006), according to the manufacturer’s instructions. PCR amplification, cloning and sequencing were performed as previously described (*Arnaud et al., 2006*). Details on the primers used can be found in Supplementary Table S1.

#### DNA methylation data mining

ES and NPC WGBS data were obtained from Geo DataSets under the accession number GSM748786 and GSM748788, respectively. The reads were first processed using TrimGalore and then mapped to the mm39 genome using Bismark. Duplicated read alignments were removed with the script deduplicate_bismark. The CpG methylation levels were computed from the selected alignments using bismark_methylation_extractor (--no_header --cutoff 4 –bedgraph) and coverage2cytosine scripts. The output was converted with the bedGraphToBigWig tools in order to be loaded on UCSC genome browser.

CpG methylation levels of ESC, NPC and frontal cortex on the mm10 genome were obtained from the tracks Stadler 2011 and Lister 2013 of the DNA methylation Hub on UCSC.

#### genomes production for NGS data alignment of hybrid samples

The sequences of the JF1 strain have been obtained from the DDBJ database under the accession number: DRP000326 and DRP000984. Paired-end reads were filtered using the CutAdapt tool to exclude poor quality reads. (--minimum-length=101 --pair-filter=any -q 20). The remaining reads were mapped on the mm39 genome with bowtie2. Alignments were filtered for poor quality with samtools wiew (-q 20). Duplicated alignments were excluded using samtools fixmate and markdup (-r). In order to identify JF1 polymorphisms, the filtered alignments were analysed using the freebayes tool (-m 20 -q 30 -C 10 -F 0.75) and the output was normalized using the tools bcftools norm. The variants were then decomposed using the vcflib vcfallelicprimitives (-kg) tool. The resulting vcf file was then processed with the mm39 genome using a custom R script in order to generate the genomes used for the alignments of various experiments (library: Biostrings, GenomicRanges). An mm39 genome masked by an N at JF1 SNP positions was generated. The JF1 genome was also reconstructed by converting JF1 SNPs, insertions and deletions in the mm39 genome. A diploid hybrid genome consisting for each chromosome of the C57BL/6 and JF1 sequences was generated. The use of the hybrid genome has the advantage of taking the JF1 indel into account when determining allelic alignments, but results in different genomic coordinates for the same element on the C57BL/6 and JF1 genomes. In order to work with only one reference, a custom R script was written to convert the coordinates of the JF1 alignments to the mm39 referential (GenomicRanges).

### Expression Analysis

#### RNA extraction

RNA was isolated from frozen cell pellets using TRIzol Reagent (Life Technologies, 15596018), according to the manufacturer’s recommendations.

#### RT-qPCR

After treatment with RNase-free DNase I (Life Technologies, 180868-015), first-strand cDNA was generated by reverse transcription with Superscript-IV (Life Technologies, 18090050) using random primers and 500 ng of RNA. cDNA was then amplified by real-time PCR with a SYBR Green mixture (Roche) using a LightCycler R 480II (Roche) apparatus.

The relative expression level was quantified with the 2-dCt method that gives the fold change variation in gene expression normalized to the geometrical mean of the expression of the housekeeping genes *Gapdh*, *Tbp* and *Gus*. The primer sequences are in Supplementary Tables S1.

##### Allelic analysis

For each locus of interest, the parental allele origin of expression was assigned following direct sequencing of the cognate RT-PCR product that encompassed a strain-specific SNP (SNP details in Supplementary Table S1).

#### Microfluidic-based quantitative analysis

This analysis was performed on a commercial panel of total RNA (RNA (mouse total RNA master panel; Ozyme 636644) obtained from pooled samples isolated from several hundreds of mouse embryos and adults. Following reverse transcription, as described above, first-strand cDNA was pre-amplified for 14 cycles with the pool of primers used for the RT-qPCR analysis and the Taq-Man PreAmplification Master Mix (Life Technologies, 4488593). RT-qPCRs were then performed and validated on Fluidigm 96.96 Dynamic Arrays using the Biomark HD system (Fluidigm Corp.) according to the manufacturer’s instructions. The relative gene expression was quantified using the 2-dCt method that gives the fold changes in gene expression normalized to the geometrical mean of the expression of the housekeeping genes *Arbp*, *Gapdh*, *Tbp*. For each condition, the presented data were obtained from two independent experiments, each analyzed in duplicate.

#### RNA-seq

Paired-end RNA-seq were generated on ES and NP cells in duplicate for the B6xJF1 and JF1xB6 genetic backgrounds. RNA-seq libraries prepared with either Illumina® TruSeq Stranded mRNA kit or NEBNext® Ultra™ II mRNA-Seq kit and sequenced on an HiSeq4000 or NovaSeq6000 apparatus were performed by Integragen SA according to the manufacturer’s protocol. To determine global and allelic expression, RNA-seq reads were mapped on the mm39 masked genome and hybrid genome, respectively, using TopHat2 and a gene annotation file adapted for these genomes based on the UCSC refGene track (-r 350 --mate-std-dev 250 --library-type fr-firststrand). Alignments were filtered with samtools for mapping quality and reads mapped in proper pair (view -f 2 -q 20). This step, on the hybrid alignments allows to isolate allele-specific mapping. The strand-specific coverages of the RNA-seq were generated on the C57BL/6 and JF1 specific alignments as well as on the global alignments using bamCoverage (--normalizeUsing RPKM –filterRNAstrand forward/ reverse) and visualized on UCSC genome browser. Replicates were overlaid for allelic and strand specific coverage using the track collection builder tool for genome exploration.

#### Gene expression data mining

Expression data of cortex at embryonic day 13.5 from B6xJF1 and JF1xB6 genetic backgrounds were obtained from GSE58523 dataset. The RNA-seq treatment was based on the pipeline describe above adapted for single end RNA-seq. Expression data of the ES cells KO for the PRC2 components were obtained from the GSE58016 dataset and were aligned to the mm39 genome and were processed for visualisation on the UCSC genome browser.

### Chromatin Immunoprecipitation Analysis

#### ChIP-qPCR

ChIP of native chromatin was performed as described by *Brind’Amour et al., 2015* using 500,000 cells per immunoprecipitation. Results presented in this article were obtained from at least three ChIP assays performed on independent chromatin preparations, as indicated in the figure legends. Details of the antisera used can be found in Supplementary Table S2. Quantitative and allelic analyses were performed as described previously (*Maupetit-Mehouas et al., 2016*). Details of the SNPs and primers used can be found in Supplementary Table 1.

#### ChIP-seq

ChiP-seq experiments on native chromatin were performed on ESC and NPC for the B6xJF1 and JF1xB6 genetic backgrounds as previously described (*Le Boiteux et al., 2021*). Details of the used antisera are in Supplementary Table S2.

Background precipitation levels were determined by performing mock precipitations with a nonspecific IgG antiserum (Sigma-Aldrich C2288), and experiments were validated by qPCR on diagnostics regions before sequencing. Library preparation (TruSeq® ChIP Sample Preparation) and sequencing on a HiSeq 2500 instrument (Illumina) were performed by MGX, according to the manufacturer’s recommendations (mean of 40 million single reads per sample). To determine global and allelic alignments, ChIP-seq reads were mapped using Bowtie2 on the mm39 masked genome and hybrid genome, respectively. Alignments filtering was realized with samtools (view -q 20), peaks were called using MACS1.4.2. (--nomodel --shiftsize 73 --pvalue 1e-5) and the coverage was computed using bamCoverage (--normalizeUsing RPKM --extendReads 200 --ignoreDuplicates --binSize 20). The track collection builder tool of UCSC was used to overlay allelic coverages for genome exploration.

#### Cut & Run

Cut&Run was performed using the CUTANA™ CUT&RUN Kit (Epicypher) on non-fixed nuclei according to the manufacturer’s instructions. Details of the antisera used are given in Supplementary Table S2. Briefly, nuclei were isolated from fresh ES or NPC cells and stored at −80°C in a nuclear extraction buffer. After thawing, 500,000 nuclei per reaction were aliquoted and incubated with pre-activated Concanavalin A-coated beads for 10 min at room temperature, followed by overnight incubation at 4°C with 0.5 ug of antibody in buffer containing 0.01% digitonin. Nuclei bound to ConA beads were then permeabilized with buffer containing 0.01% digitonin and incubated with pAG-MNase fusion protein for 10 min at room temperature. After washing, cleavage of chromatin-bound pAG-MNase was induced by the addition of calcium chloride to a final concentration of 2mM. After incubation at 4°C for 2 h, the reaction was stopped by the addition of STOP buffer (containing fragmented genomic E. coli DNA as spike-in). Following fragmented DNA purification, Illumina sequencing libraries were prepared from ∼5 ng of purified DNA using the CUTANA™ CUT&RUN Library Prep Kit (EpiCypher 14-1001 & 14-1002) according to the manufacturer’s recommendations. Purified multiplex libraries were diluted to 9 nM concentration (calculated based on Qubit dsDNA HS Assay Kit) and sequenced on a NovaSeq 6000 instrument (Illumina) from IntegraGen SA. Paired-end reads were mapped to the mm39 masked genome and hybrid genome using Bowtie2. Alignments filtering was realized with samtools (view -f 2 -q 20), and the coverage was obtained using bamCoverage (for the global coverage: --scaleFactor “spike-in DNA” --normalizeUsing RPKM --binSize 25, for the allelic coverage: --binSize 25). The track collection builder tool of UCSC was used to overlay allelic coverages for genome exploration. Peaks were called with MACS2 using Cut&Run sample performed with an IgG as a control (callpeak -f BAMPE --keep-dup all).

#### ChIP-seq and ATAC -seq data mining

Allelic and global mm9 alignments for H3K27ac in frontal cortex were obtained from the datasets GSM751461 and GSM751462. These alignments were converted into coverages using a R script. (rtracklayer, GenomicRanges) and were visualized on UCSC.

ATAC-seq data for ES and ES derived neural progenitor cells were obtained from Geo DataSets GSE155215. Paired-end reads were treated with trim_galore (--paired) and then were aligned on the mm39 genome using bowtie2 (--very-sensitive -X 1000). Only proper paired alignments were conserved with samtools (view -f 2) and alignments on mitochondrial and random chromosomes were excluded. PCR duplicates were removed using picard-tools (MarkDuplicates --REMOVE_DUPLICATES=true). The coverage was produced using bamCoverage (--normalizeUsing RPKM --binSize 20) and was visualized on UCSC genome browser. Peaks were called using macs2 (callpeak -f BAMPE --broad --broad-cutoff 0.05 --keep-dup all).

### 4C-seq

4C-seq experiments were generated on ES and NP cells from B6xJF1 and JF1xB6 genetic backgrounds for various viewpoints. Details of the primers used for each viewpoint can be found in Supplementary Table 1. 4C Template Preparation was carried out as previously described with some modifications (*Harmen J G van de Werken et al,).* Briefly, 1×107 cell suspensions were cross-linked with formaldehyde (final concentration 2%) for 10 min. After cell lysis and permeabilization by SDS and Triton X-100 treatments, samples were digested with 600U DpnII at 37°C in 1X NEBuffer DpnII (4 h with 200U, overnight with 200U and 4 h with 200U). The restriction enzyme was inactivated with SDS and Triton X-100. The first ligation was performed overnight at 18°C in a large volume, 7.2 ml of 1X ligase buffer and with 50U T4 DNA Ligase. The cross-linking was reversed with 600ug Proteinase K at 65°C overnight. After phenol/chloroform purification, the DNA was digested with 50U of NlaIII overnight at 37°C. After phenol/chloroform purification, a second ligation was performed overnight at 18°C in 14ml of 1X ligase buffer and with 100U of T4 DNA Ligase. The 4C Template was concentrated with an ethanol precipitation and then purified using the DNA Clean & Concentrator Zymo-25 kit. To produce a 4C-seq library, 3,2 µg of 4C template was amplified into 16 PCR reactions with viewpoint specific sequencing primers and 56 U of Expand Long Template Polymerase. PCR reactions were pooled and purified using High Pure PCR Product Purification Kit. 4C-seq library of different viewpoints were combined before sequencing on an HiSeq 4000 or NovaSeq 6000 instrument (Illumina) by IntegraGen SA. Due to the primer design, paired end reads were used to determine the viewpoint allele and the interacting sequence. To do this, only the expected sequence, corresponding to the viewpoint of the informative reads, was mapped to the mm39 hybrid genome using bowtie2 (--trim5 10 --trim3 “viewpoint specific” --local --very-sensitive-local). Only alignments mapped to the viewpoint coordinates and with a minimal quality were conserved to determine the origin allelic of the viewpoint in the reads (samtools view -q 10 [viewpoint coordinates]). To determine the sequence in interaction, the expected sequence with DpnII restriction site was mapped to the mm39 masked genome using bowtie2 (--trim5 “viewpoint specific” --trim3 “viewpoint specific” --local --very-sensitive-local). these alignments were filtered for mapping quality (samtools view -q 10) and were split according to the origin allelic of the viewpoint. To construct the allelic interactome, the alignments were processed the FourCSeq Bioconductor package to count the reads mapped exactly to the end of a DpnII fragment and to generate a smoothed rpm normalized coverage. The track collection builder tool of UCSC was used to overlay allelic interactome coverages for genome exploration.

### Reanalysis of HI-C

HI-C data were obtained from Geo DataSets under the accession numbers: for ES cells (GSM2533818, GSM2533819, GSM2533820, GSM2533821) and for in vivo NPC (GSM2533835, GSM2533836, GSM2533837, GSM2533838). These paired end reads were processed with HiC-Pro (*Servant N et al,* Genome Biology (2015), 16:259) with the following parameters (MIN_MAPQ = 20, REFERENCE_GENOME = mm39, GENOME_FRAGMENT = DpnII_resfrag_hg19.bed, LIGATION_SITE = GATC, BIN_SIZE = 5000). The produced normalized contact matrix (iced) was used to generate the contact map with the HiTC Bioconductor package.

## Reference

Sondheimer, N., and Lindquist, S. (2000). Rnq1: an epigenetic modifier of protein function in yeast. Mol. Cell 5, 163–172. 10.1016/S1097-2765(00)80412-8.

Andergassen D, Dotter CP, Wenzel D, Sigl V, Bammer PC, Muckenhuber M, Mayer D, Kulinski TM, Theussl HC, Penninger JM, Bock C, Barlow DP, Pauler FM, Hudson QJ. (2017) Mapping the mouse Allelome reveals tissue-specific regulation of allelic expression. Elife 6:e25125. doi: 10.7554/eLife.25125.

Arnaud P, Hata K, Kaneda M, Li E, Sasaki H, Feil R, Kelsey G. (2006) Stochastic imprinting in the progeny of Dnmt3L-/- females. Hum Mol Genet. 15(4):589–98. doi: 10.1093/hmg/ddi475.

Aslanger AD, Goncu B, Duzenli OF, Yucesan E, Sengenc E, Yesil G. (2022) Biallelic loss of TRAPPC9 function links vesicle trafficking pathway to autosomal recessive intellectual disability. J Hum Genet. 67(5):279–284. doi: 10.1038/s10038-021-01007-8.

Babak T, DeVeale B, Tsang EK, Zhou Y, Li X, Smith KS, Kukurba KR, Zhang R, Li JB, van der Kooy D, Montgomery SB, Fraser HB. (2015) Genetic conflict reflected in tissue-specific maps of genomic imprinting in human and mouse. Nat Genet. 47(5):544–9. doi: 10.1038/ng.3274.

Bartman CR, Hsu SC, Hsiung CC, Raj A, Blobel GA. (2016) Enhancer Regulation of Transcriptional Bursting Parameters Revealed by Forced Chromatin Looping. Mol Cell. 62(2):237–247. doi: 10.1016/j.molcel.2016.03.007.

Bonev B, Mendelson Cohen N, Szabo Q, Fritsch L, Papadopoulos GL, Lubling Y, Xu X, Lv X, Hugnot JP, Tanay A, Cavalli G. (2017) Multiscale 3D Genome Rewiring during Mouse Neural Development. Cell. 171(3):557–572.e24. doi: 10.1016/j.cell.2017.09.043.

Bonthuis PJ, Huang WC, Stacher Hörndli CN, Ferris E, Cheng T, Gregg C. (2015) Noncanonical Genomic Imprinting Effects in Offspring. Cell Rep. 12(6):979–91. doi: 10.1016/j.celrep.2015.07.017.

Bouschet T, Dubois E, Reynès C, Kota SK, Rialle S, Maupetit-Méhouas S, Pezet M, Le Digarcher A, Nidelet S, Demolombe V, Cavelier P, Meusnier C, Maurizy C, Sabatier R, Feil R, Arnaud P, Journot L, Varrault A. (2017) In Vitro Corticogenesis from Embryonic Stem Cells Recapitulates the In Vivo Epigenetic Control of Imprinted Gene Expression. Cereb Cortex. 27(3):2418–2433. doi: 10.1093/cercor/bhw102.

Brind’Amour J, Liu S, Hudson M, Chen C, Karimi MM, Lorincz MC. (2015) An ultra-low-input native ChIP-seq protocol for genome-wide profiling of rare cell populations. Nat Commun. 6:6033. doi: 10.1038/ncomms7033.

Claxton M, Pulix M, Seah MKY, Bernardo R, Zhou P, Aljuraysi S, Liloglou T, Arnaud P, Kelsey G, Messerschmidt DM, Plagge A. (2022) Variable allelic expression of imprinted genes at the *Peg13*, *Trappc9*, Ago2 cluster in single neural cells. Front Cell Dev Biol. 10:1022422. doi: 10.3389/fcell.2022.1022422.

Court F, Baniol M, Hagege H, Petit JS, Lelay-Taha MN, Carbonell F, Weber M, Cathala G, Forne T. (2011) Long-range chromatin interactions at the mouse Igf2/H19 locus reveal a novel paternally expressed long non-coding RNA. Nucleic Acids Res. 39:5893–906.

Court F, Camprubi C, Garcia CV, Guillaumet-Adkins A, Sparago A, Seruggia D, Sandoval J, Esteller M, Martin-Trujillo A, Riccio A, Montoliu L, Monk D. (2014) The PEG13-DMR and brain-specific enhancers dictate imprinted expression within the 8q24 intellectual disability risk locus. Epigenetics Chromatin. 7:5.

Cho SW, Xu J, Sun R, Mumbach MR, Carter AC, Chen YG, Yost KE, Kim J, He J, Nevins SA, Chin SF, Caldas C, Liu SJ, Horlbeck MA, Lim DA, Weissman JS, Curtis C, Chang HY. (2018) Promoter of lncRNA Gene PVT1 Is a Tumor-Suppressor DNA Boundary Element. Cell. 173(6):1398–1412.e22. doi: 10.1016/j.cell.2018.03.068.

De Gobbi M, Viprakasit V, Hughes JR, Fisher C, Buckle VJ, Ayyub H, Gibbons RJ, Vernimmen D, Yoshinaga Y, de Jong P, Cheng JF, Rubin EM, Wood WG, Bowden D, Higgs DR. (2006) A regulatory SNP causes a human genetic disease by creating a new transcriptional promoter. Science. 312(5777):1215–7. doi: 10.1126/science.1126431.

Dixon JR, Selvaraj S, Yue F, Kim A, Li Y, Shen Y, Hu M, Liu JS, Ren B. (2012) Topological domains in mammalian genomes identified by analysis of chromatin interactions. Nature. 485:376–80.

Gaspard N, Bouschet T, Herpoel A, Naeije G, van den Ameele J, Vanderhaeghen P (2009) Generation of cortical neurons from mouse embryonic stem cells. Nat Protoc 4:1454–1463

Hanna CW, Kelsey G. (2021) Features and mechanisms of canonical and noncanonical genomic imprinting. Genes Dev. 35(11-12):821–834. doi: 10.1101/gad.348422.121.

Ghavi-Helm Y, Klein FA, Pakozdi T, Ciglar L, Noordermeer D, Huber W, Furlong EE. (2014) Enhancer loops appear stable during development and are associated with paused polymerase. Nature. 512(7512):96–100. doi: 10.1038/nature13417.

Hata K, Okano M, Lei H, Li E. (2002) Dnmt3L cooperates with the Dnmt3 family of de novo DNA methyltransferases to establish maternal imprints in mice. Development 129(8):1983–93. doi: 10.1242/dev.129.8.1983.

Henckel A, Arnaud P. (2010) Genome-wide identification of new imprinted genes. Brief Funct Genomics 304–14. doi: 10.1093/bfgp/elq016.

Hosogane M, Funayama R, Shirota M, Nakayama K. (2016) Lack of Transcription Triggers H3K27me3 Accumulation in the Gene Body. Cell Rep. 16(3):696–706. doi: 10.1016/j.celrep.2016.06.034.

Huang D, Ovcharenko I. (2022) Enhancer-silencer transitions in the human genome. Genome Res. 32(3):437–448. doi: 10.1101/gr.275992.121.

Ivanova E, Kelsey G. (2011) Imprinted genes and hypothalamic function. J Mol Endocrinol. 47:R6774.

Juan AM, Foong YH, Thorvaldsen JL, Lan Y, Leu NA, Rurik JG, Li L, Krapp C, Rosier CL, Epstein JA, Bartolomei MS. (2022) Tissue-specific Grb10/Ddc insulator drives allelic architecture for cardiac development. Mol Cell. 82(19):3613–3631.e7. doi: 10.1016/j.molcel.2022.08.021.

Kim TK, Hemberg M, Gray JM, Costa AM, Bear DM, Wu J, Harmin DA, Laptewicz M, Barbara-Haley K, Kuersten S, Markenscoff-Papadimitriou E, Kuhl D, Bito H, Worley PF, Kreiman G, Greenberg ME. (2010) Widespread transcription at neuronal activity-regulated enhancers. Nature 465(7295):182–7. doi: 10.1038/nature09033.

Kurukuti S, Tiwari VK, Tavoosidana G, Pugacheva E, Murrell A, Zhao Z, Lobanenkov V, Reik W, Ohlsson R. (2006) CTCF binding at the H19 imprinting control region mediates maternally inherited higher-order chromatin conformation to restrict enhancer access to Igf2. Proc Natl Acad Sci U S A;103(28):10684–9. doi: 10.1073/pnas.0600326103.

Lau JC, Hanel ML, Wevrick R (2004) Tissue-specific and imprinted epigenetic modifications of the human NDN gene. Nucleic Acids Res 32: 3376–3382

Le Boiteux E, Court F, Guichet PO, Vaurs-Barrière C, Vaillant I, Chautard E, Verrelle P, Costa BM, Karayan-Tapon L, Fogli A, Arnaud P. (2021) Widespread overexpression from the four DNA hypermethylated HOX clusters in aggressive (IDHwt) glioma is associated with H3K27me3 depletion and alternative promoter usage. Mol Oncol.15(8)1995–2010. doi: 10.1002/1878-0261.12944.

Lessel D, Zeitler DM, Reijnders MRF, Kazantsev A, Hassani Nia F, Bartholomäus A, Martens V, Bruckmann A, Graus V, McConkie-Rosell A, McDonald M, Lozic B, Tan ES, Gerkes E, Johannsen J, Denecke J, Telegrafi A, Zonneveld-Huijssoon E, Lemmink HH, Cham BWM, Kovacevic T, Ramsdell L, Foss K, Le Duc D, Mitter D, Syrbe S, Merkenschlager A, Sinnema M, Panis B, Lazier J, Osmond M, Hartley T, Mortreux J, Busa T, Missirian C, Prasun P, Lüttgen S, Mannucci I, Lessel I, Schob C, Kindler S, Pappas J, Rabin R, Willemsen M, Gardeitchik T, Löhner K, Rump P, Dias KR, Evans CA, Andrews PI, Roscioli T, Brunner HG, Chijiwa C, Lewis MES, Jamra RA, Dyment DA, Boycott KM, Stegmann APA, Kubisch C, Tan EC, Mirzaa GM, McWalter K, Kleefstra T, Pfundt R, Ignatova Z, Meister G, Kreienkamp HJ. (2020) Germline AGO2 mutations impair RNA interference and human neurological development. Nat Commun.11(1):5797. doi: 10.1038/s41467-020-19572-5.

Li T, Vu TH, Ulaner GA, Yang Y, Hu JF, Hoffman AR (2004) Activating and silencing histone modifications form independent allelic switch regions in the imprinted Gnas gene. Hum Mol Genet13: 741–750

Liang ZS, Cimino I, Yalcin B, Raghupathy N, Vancollie VE, Ibarra-Soria X, Firth HV, Rimmington D, Farooqi IS, Lelliott CJ, Munger SC, O’Rahilly S, Ferguson-Smith AC, Coll AP, Logan DW. Trappc9 deficiency causes parent-of-origin dependent microcephaly and obesity. PLoS Genet. 2020 Sep 2;16(9):e1008916. doi: 10.1371/journal.pgen.1008916.

Lin S, Ferguson-Smith AC, Schultz RM, Bartolomei MS. (2011) Nonallelic transcriptional roles of CTCF and cohesins at imprinted loci. Mol Cell Biol. 31(15):3094–104. doi: 10.1128/MCB.01449-10. Epub 2011 May 31.

Llères D, Moindrot B, Pathak R, Piras V, Matelot M, Pignard B, Marchand A, Poncelet M, Perrin A, Tellier V, Feil R, Noordermeer D. (2019) CTCF modulates allele-specific sub-TAD organization and imprinted gene activity at the mouse Dlk1-Dio3 and Igf2-H19 domains. Genome Biol. 20(1):272. doi: 10.1186/s13059-019-1896-8.

Mager J, Montgomery ND, de Villena FP, Magnuson T (2003) Genome imprinting regulated by the mouse Polycomb group protein Eed. Nat Genet 33: 502–507

Maupetit-Méhouas S, Montibus B, Nury D, Tayama C, Wassef M, Kota SK, Fogli A, Cerqueira Campos F, Hata K, Feil R, Margueron R, Nakabayashi K, Court F, Arnaud P. (2016) Imprinting control regions (ICRs) are marked by mono-allelic bivalent chromatin when transcriptionally inactive. Nucleic Acids Res. 44:621–35. doi: 10.1093/nar/gkv960.

Monk D, Arnaud P, Apostolidou S, Hills FA, Kelsey G, Stanier P, Feil R, Moore GE. (2006) Limited evolutionary conservation of imprinting in the human placenta. Proc Natl Acad Sci U S A. 103(17):6623–8. doi: 10.1073/pnas.0511031103.

Montibus B, Cercy J, Bouschet T, Charras A, Maupetit-Méhouas S, Nury D, Gonthier-Guéret C, Chauveau S, Allegre N, Chariau C, Hong CC, Vaillant I, Marques CJ, Court F, Arnaud P. (2021) TET3 controls the expression of the H3K27me3 demethylase Kdm6b during neural commitment. Cell Mol Life Sci. 78(2):757–768. doi: 10.1007/s00018-020-03541-8.

Murrell A, Heeson S, Reik W. (2004) Interaction between differentially methylated regions partitions the imprinted genes Igf2 and H19 into parent-specific chromatin loops. Nat Genet. 36(8):889–93.

Neely MD, Litt MJ, Tidball AM, Li GG, Aboud AA, Hopkins CR, Chamberlin R, Hong CC, Ess KC, Bowman AB (2012) DMH1, a highly selective small molecule BMP inhibitor promotes neurogenesis of hiPSCs: comparison of PAX6 and SOX1 expression during neural induction. ACS Chem Neurosci 3:482–491.

Noordermeer D, Feil R. (2020) Differential 3D chromatin organization and gene activity in genomic imprinting. Curr Opin Genet Dev. 61:17–24. doi: 10.1016/j.gde.2020.03.004.

Oudelaar AM, Harrold CL, Hanssen LLP, Telenius JM, Higgs DR, Hughes JR. (2019) A revised model for promoter competition based on multi-way chromatin interactions at the α-globin locus. Nat Commun. 10(1):5412. doi: 10.1038/s41467-019-13404-x.

Pauler FM, Sloane MA, Huang R, Regha K, Koerner MV, Tamir I, Sommer A, Aszodi A, Jenuwein T, Barlow DP. (2008) H3K27me3 forms BLOCs over silent genes and intergenic regions and specifies a histone banding pattern on a mouse autosomal chromosome. Genome Res.19(2):221–33. doi: 10.1101/gr.080861.108.

Prickett AR, Barkas N, McCole RB, Hughes S, Amante SM, Schulz R, Oakey RJ. (2013) Genome-wide and parental allele-specific analysis of CTCF and cohesin DNA binding in mouse brain reveals a tissue-specific binding pattern and an association with imprinted differentially methylated regions. Genome Res. 23(10):1624–35.

Ruf N, Bähring S, Galetzka D, Pliushch G, Luft FC, Nürnberg P, Haaf T, Kelsey G, Zechner U. (2007) Sequence-based bioinformatic prediction and QUASEP identify genomic imprinting of the KCNK9 potassium channel gene in mouse and human. Hum Mol Genet. 16(21):2591–9. doi: 10.1093/hmg/ddm216.

Shukla V, Cetnarowska A, Hyldahl M, Mandrup S. (2022) Interplay between regulatory elements and chromatin topology in cellular lineage determination. Trends Genet. 38(10):1048–1061. doi: 10.1016/j.tig.2022.05.011.

Simeoni F, Romero-Camarero I, Camera F, Amaral FMR, Sinclair OJ, Papachristou EK, Spencer GJ, Lie-A-Ling M, Lacaud G, Wiseman DH, Carroll JS, Somervaille TCP. (2021) Enhancer recruitment of transcription repressors RUNX1 and TLE3 by mis-expressed FOXC1 blocks differentiation in acute myeloid leukemia. Cell Rep. 36(12):109725.

Smith RJ, Dean W, Konfortova G, Kelsey G. (2003) Identification of novel imprinted genes in a genome-wide screen for maternal methylation. Genome Res. 13(4):558–69. doi: 10.1101/gr.781503.

Topfer SK, Feng R, Huang P, Ly LC, Martyn GE, Blobel GA, Weiss MJ, Quinlan KGR, Crossley M. (2022) Disrupting the adult globin promoter alleviates promoter competition and reactivates fetal globin gene expression. Blood 139(14):2107–2118. doi: 10.1182/blood.2021014205.

Tucci V, Isles AR, Kelsey G, Ferguson-Smith AC; Erice Imprinting Group. (2019) Genomic Imprinting and Physiological Processes in Mammals. Cell 176(5):952–965. doi: 10.1016/j.cell.2019.01.043.

Umlauf, D., Goto, Y., Cao, R., Cerqueira, F., Wagschal, A., Zhang, Y. and Feil, R. (2004) Imprinting along the Kcnq1 domain on mouse chromosome 7 involves repressive histone methylation and recruitment of Polycomb group complexes. Nat. Genet., 36, 1296–1300.

van de Werken HJ, de Vree PJ, Splinter E, Holwerda SJ, Klous P, de Wit E, de Laat W. (2012) 4C technology: protocols and data analysis. Methods Enzymol. 513:89–112. doi: 10.1016/B978-0-12-391938-0.00004-5.

Visel A, Minovitsky S, Dubchak I, Pennacchio LA (2007). VISTA Enhancer Browser-a database of tissue-specific human enhancers. Nucleic Acids Res 35:D88–92

Wagschal A, Sutherland HG, Woodfine K, Henckel A, Chebli K, Schulz R, Oakey RJ, Bickmore WA, Feil R (2008) G9a histone methyltransferase contributes to imprinting in the mouse placenta. Mol Cell Biol 28: 1104–1113

Wilton KM, Gunderson LB, Hasadsri L, Wood CP, Schimmenti LA. (2020) Profound intellectual disability caused by homozygous TRAPPC9 pathogenic variant in a man from Malta. Mol Genet Genomic Med. 8(5):e1211. doi: 10.1002/mgg3.1211.

Wurmser A, Basu S. (2022) Enhancer-Promoter Communication: It’s Not Just About Contact. Front Mol Biosci. 9:867303. doi: 10.3389/fmolb.2022.867303.

Xiao JY, Hafner A, Boettiger AN. (2021) How subtle changes in 3D structure can create large changes in transcription. Elife 10:e64320. doi: 10.7554/eLife.64320.

Yang Y, Li T, Vu TH, Ulaner GA, Hu JF, Hoffman AR (2003) The histone code regulating expression of the imprinted mouse Igf2r gene. Endocrinology 144: 5658–5670

Young MD, Willson TA, Wakefield MJ, Trounson E, Hilton DJ, Blewitt ME, Oshlack A, Majewski IJ. (2011) ChIP-seq analysis reveals distinct H3K27me3 profiles that correlate with transcriptional activity. Nucleic Acids Res. 39(17):7415–27. doi: 10.1093/nar/gkr416.

Zhang S, Übelmesser N, Barbieri M, Papantonis A. (2023) Enhancer-promoter contact formation requires RNAPII and antagonizes loop extrusion. Nat Genet. 55(5):832–840. doi: 10.1038/s41588-023-01364-4.

