## Supplementary Figures for "Biallelic non-productive enhancer-promoter interaction precedes imprinted expression of *Kcnk9* during mouse neural commitment"

**Supp Figure 1: Expression patterns at *Peg13* domain in the corticogenesis model recapitulate those observed in the embryonic brain in vivo**

**A)** Microfluidic-based RT-qPCR analysis of *Kcnk9* and *Peg13* in the indicated tissues and cell types. Results are presented as the fold enrichment of the mean expression level detected in all tissues, after normalization to the geometric mean of the expression of the three housekeeping genes *Arbp*, *Gapdh* and *Tbp*. Data were obtained from two independent experiments, each analysed in duplicate. **B)** P*ou5f1*, *Nestin* and *Pax6* expression levels in ES cells (n=6) and in NP cells at day 12 (n=6) of in vitro corticogenesis. Statistical significance was determined with the unpaired t test (p values in the figure). **C)** Quantitative RT-PCR analyses to assess the expression levels of *Trappc9, Chrac1* and *Ago2* in ES cells (*n* = 4) and at day 4 (D4; *n* = 2), D6 (n=2), D8 (*n* = 2), and NPC (D12; *n* = 4) stages of in vitro corticogenesis. The parental origin of expression is shown on the lower panel. **D)** Genome browser view at the *Peg13* domain to show the allelic oriented RNA-seq signal in ESC, NPC and embryonic brain (Dorsal Telencephalon ((DT); reanalysed data from *Bouschet et al., 2016*). For each condition the quantitative and the merged allelic RNA-seq signals are shown in the upper and lower-panel, respectively. Maternal and paternal expression are shown in red and blue respectively. In **B)** to **D)** results are presented as the percentage of expression relative to the geometric mean of the expression of the three housekeeping genes *Gapdh*, *Gus* and *Tbp*. The data are presented as the mean ± SEM.

**Supp Figure 2: *Peg13 DMR* lost methylation in E9.5 *Dnmt3l* -/+ embryo**

Map of the mouse *Peg13* locus showing the CpG islands (CGI) as described in the UCSC Genome Browser. The bisulfite analysed region is shown in purple. The lower panel shows bisulfite-derived data from one WT and two Dnmt3L$^{-/+}$ embryos. Each horizontal row of circles represents the CpG dinucleotides on a single chromosome. Solid circles, methylated CpG dinucleotides; open circles, unmethylated CpG dinucleotides. Parental origin (Mat., maternal; Pat., paternal) was determined using strain-specific SNPs. Red rectangles indicate CpGs that are missing due to SNPs.

**Supp Figure 3: *Peg13* DMR and *Kcnk9* methylation patterns**

**A)** Map of the mouse *Peg13* locus showing the CpG islands (CGI) as described in the UCSC Genome Browser. The bisulfite analysed region is shown in purple. The lower panel shows bisulfite derived data from ESC and NPC, respectively. Each horizontal row of circles represents the CpG dinucleotides on a single chromosome. Solid circles, methylated CpG dinucleotides; open circles, unmethylated CpG dinucleotides. Parental origin (Mat., maternal; Pat., paternal) was determined using strain-specific SNPs. Red rectangles indicate CpGs that are missing due to SNPs. **B)** Genome Browser view at the *Kcnk9* locus to show CpG island (CGI) position and non-allelic WGBS methylation dataset from mouse ESC, NPC and frontal cortex.

**Supplementary Figure 4: Similar parental allelic 4C-seq signals for *Peg13* DMR viewpoint in reciprocal B6/JF1 and JF1/Bl6 ESC.**

Genome browser view at the *Kcnk9 -Trappc9* region to show B6/JF1 (upper panel) and JF1/Bl6 (lower panel) ESC allelic 4C-seq, for the *Peg13* DMR viewpoint, and CTCF C&R signals. 4C-seq are shown by merging the allelic signals, contacts mediated by the paternal and maternal alleles are shown in blue and red, respectively. The ratio of maternal/paternal interactions is indicated. The relative position of TAD and the imprinted domain is shown. *Peg13* DMR interactions are largely confined to the imprinted domain, within the TAD.

**Supp Figure 5: *Peg13* DMR and *PE* interactomes structure the higher-order chromatin conformation at the *Peg13* domain**

Genome Browser view at the *Kcnk9 -Trappc9* region to show in ESC (upper panel) and NPC (bottom panel); re-analysed HiC data, allelic 4C-seq for the *Peg13* DMR, *Kcnk9* promoter and the PE viewpoints and and CTCF C&R signals. 4C-seq are presented by merging the allelic signals, contacts mediated by the paternal and maternal alleles are shown in blue and red, respectively. The ratio of maternal/paternal interactions is indicated. The sub-TAD containing the *Kcnk9-Peg13* region is delineated by a dotted line. a and b denote the contact between PE and the *Kcnk9* promoter and the intronic regions, respectively.

**Supp Figure 6: *Peg13* DMR and *PE* interactomes structure the higher-order chromatin conformation at the *Peg13* domain**

**A)** Genome Browser view at the *PE* region (highlighted in grey) to show non-allelic WGBS methylation dataset from mouse ESC, NPC and frontal cortex. **B)** and **C)** Quantitative RT-PCR analyses to assess PE-associated eRNA expression in neonatal brains (n=4) in **B)** adult brains (n=2) and **C)** in ES cells (*n* =3) and at day 4 (D4; *n* = 2), D6 (n=2), D8 (*n* = 2), and NPC (D12; *n* = 3) stages of in vitro corticogenesis. Statistical significance was determined with the unpaired t test (p values in the figure). The data are presented as the mean ± SEM. The parental origin of expression is shown on the lower panel. **D)** Genome Browser view at the Peg13 and *PE* region, respectively, to show allelic H3K27ac dataset from adult mouse frontal cortex.

### Figure S1.

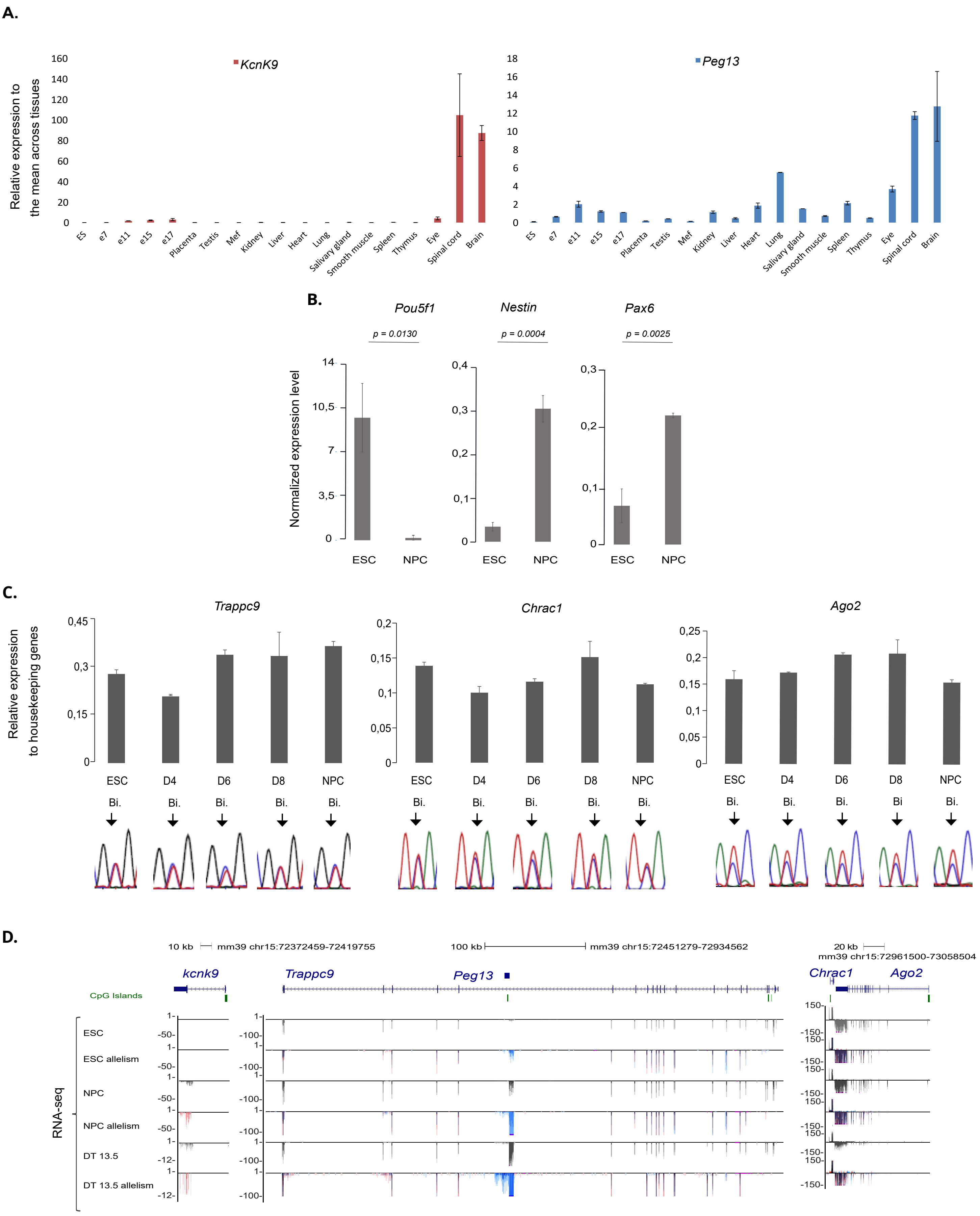
A.
KcnK9
Peg13
Relative expression to the mean across tissues
ES
e7
e11
e15
e17
Placenta
Testis
Mef
Kidney
Liver
Heart
Lung
Salivary gland
Smooth muscle
Spleen
Thymus
Eye
Spinal cord
Brain
B.
Pou5f1
Nestin
Pax6
p = 0.0130
p = 0.0004
p = 0.0025
Normalized expression level
ESC
NPC
C.
Trappc9
Chrac1
Ago2
Relative expression to housekeeping genes
ESC
D4
D6
D8
NPC
Bi.
D.
10 kb
mm39 chr15:72372459-72419755
100 kb
mm39 chr15:72451279-72934562
20 kb
mm39 chr15:72961500-73058504
kcnk9
Trappc9
Peg13
Chrac1
Ago2
CpG Islands
RNA-seq
ESC
ESC allelism
NPC
NPC allelism
DT 13.5
DT 13.5 allelism

**Figure S2.**

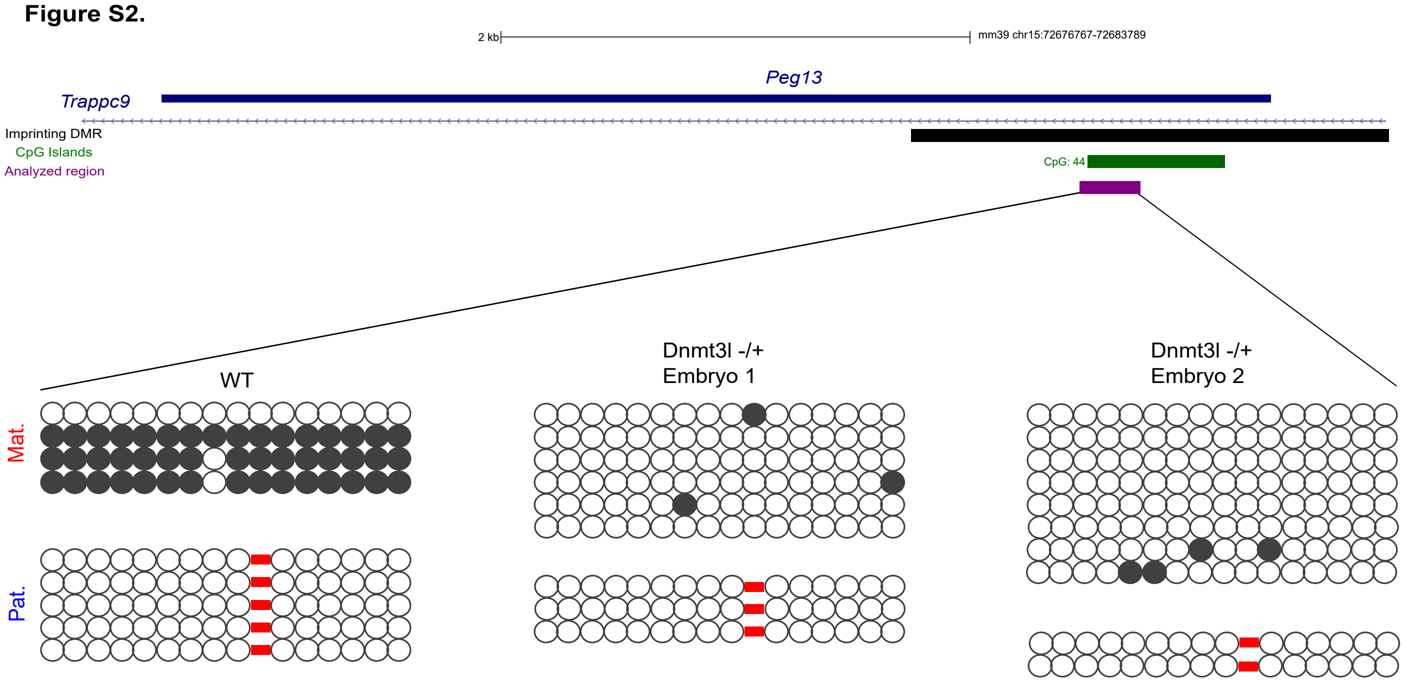
2 kb
mm39 chr15:72676767-72683789
Peg13
Trappc9
Imprinting DMR
CpG Islands
CpG: 44
Analyzed region
WT
Dnmt3l -/+
Embryo 1
Dnmt3l -/+
Embryo 2
Mat.
Pat.

**Figure S3.**

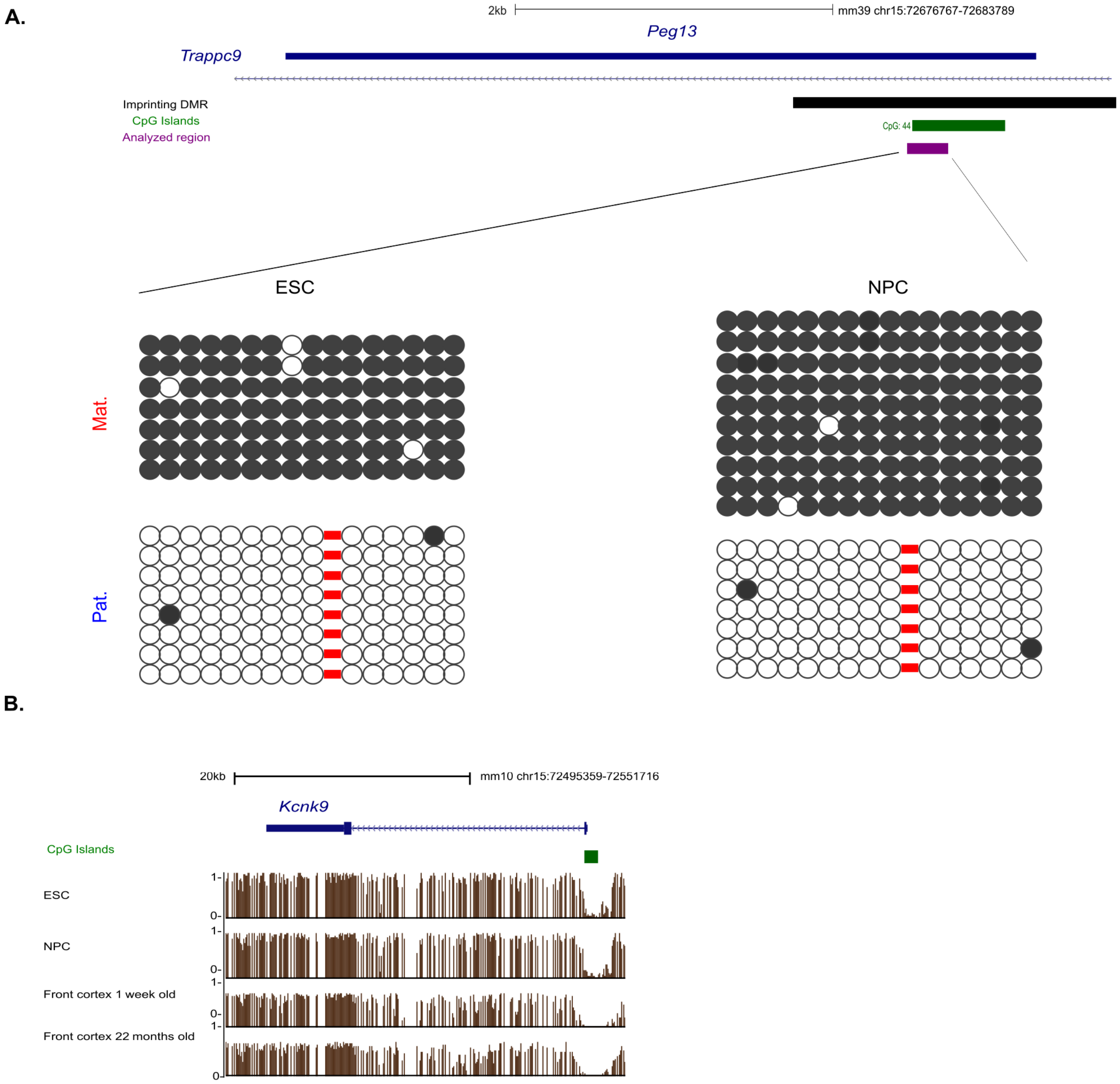

A.
2kb
mm39 chr15:72676767-72683789
Peg13
Trappc9
Imprinting DMR
CpG Islands
Analyzed region
CpG: 44
ESC
NPC
Mat.
Pat.
B.
20kb
mm10 chr15:72495359-72551716
Kcnk9
CpG Islands
ESC
NPC
Front cortex 1 week old
Front cortex 22 months old
1-
0-

**Figure S4.**

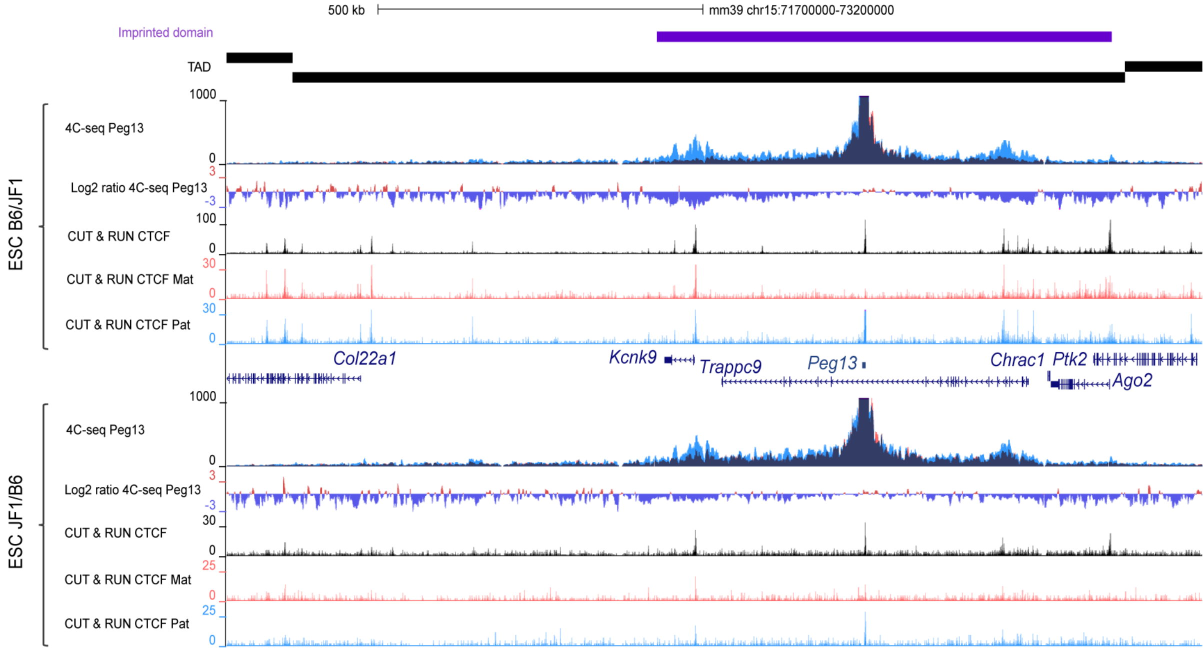

500 kb
mm39 chr15:71700000-73200000
Imprinted domain
TAD
ESC B6/JF1
1000
4C-seq Peg13
0
3
Log2 ratio 4C-seq Peg13
-3
100
CUT & RUN CTCF
0
30
CUT & RUN CTCF Mat
0
30
CUT & RUN CTCF Pat
0
Col22a1
Kcnk9
Trappc9
Peg13
Chrac1
Ptk2
Ago2
ESC JF1/B6
1000
4C-seq Peg13
0
3
Log2 ratio 4C-seq Peg13
-3
30
CUT & RUN CTCF
0
25
CUT & RUN CTCF Mat
0
25
CUT & RUN CTCF Pat
0

**Figure S5.**

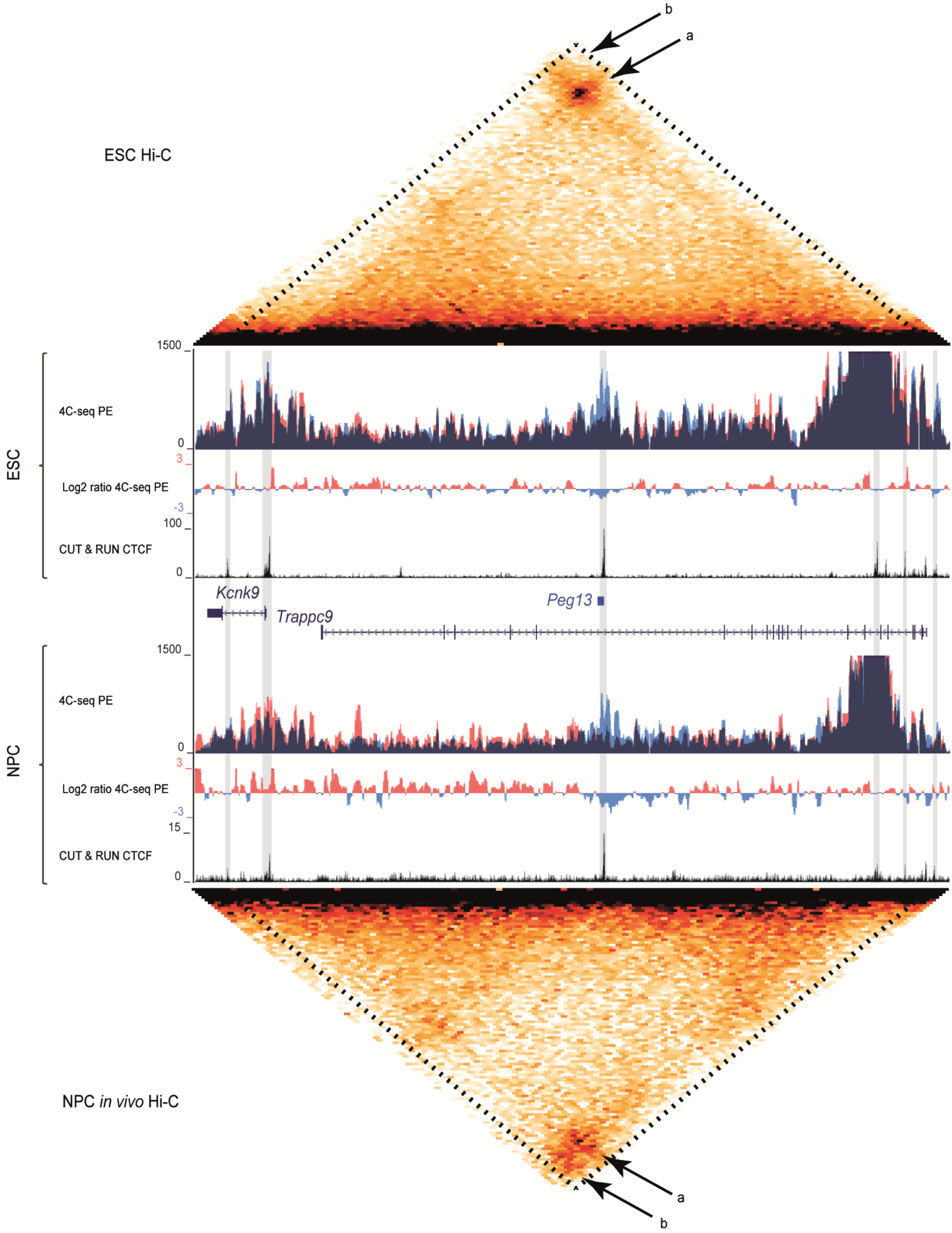

b
a
ESC Hi-C
1500
4C-seq PE
0
3
Log2 ratio 4C-seq PE
-3
100
CUT & RUN CTCF
0
ESC
Kcnk9
Peg13
Trappc9
1500
4C-seq PE
0
3
Log2 ratio 4C-seq PE
-3
15
CUT & RUN CTCF
0
NPC
NPC in vivo Hi-C
a
b

**Figure S6.**

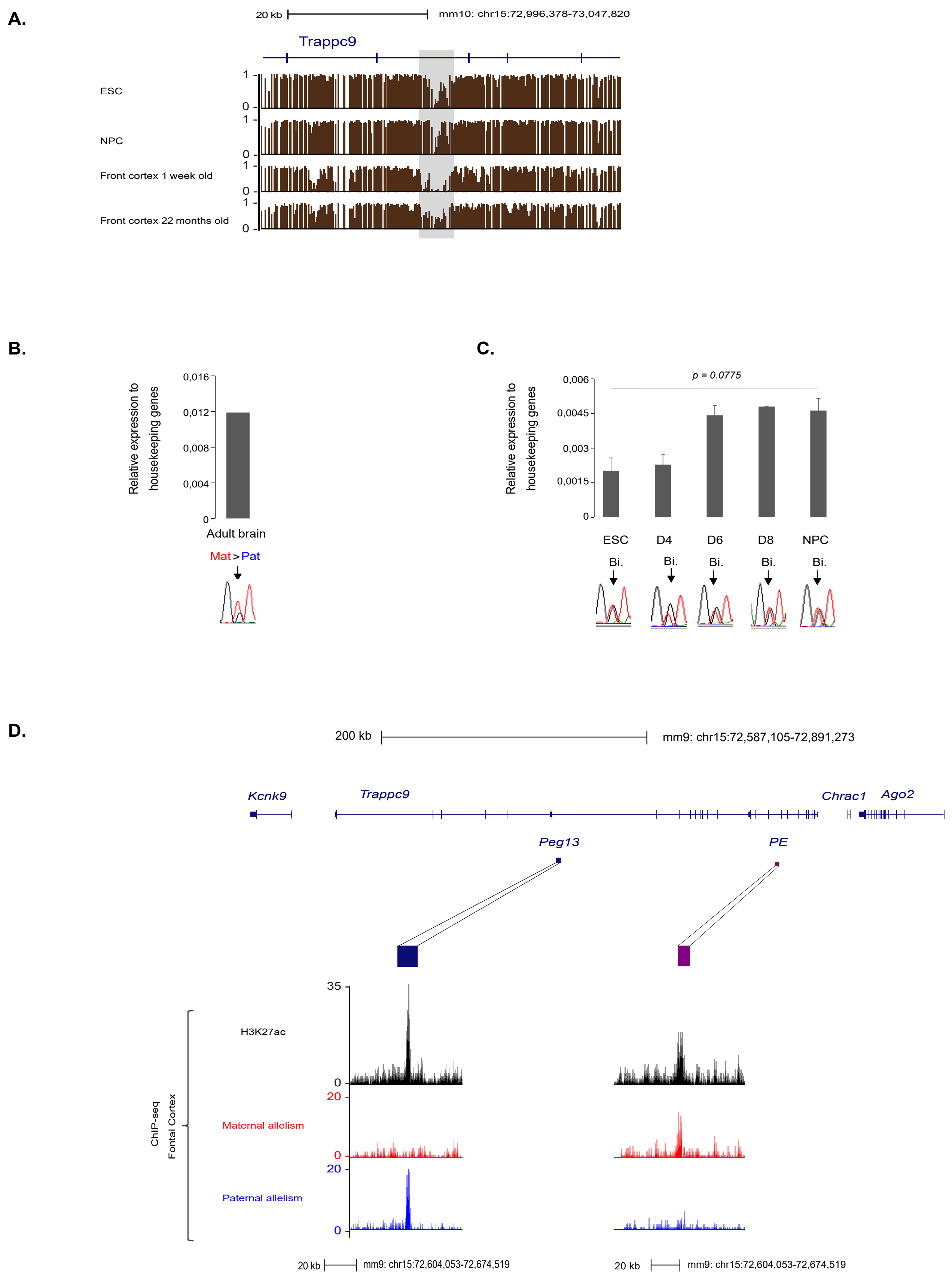
A.
20 kb
mm10: chr15:72,996,378-73,047,820
Trappc9
ESC
NPC
Front cortex 1 week old
Front cortex 22 months old
1
0
B.
Relative expression to housekeeping genes
0,016
0,012
0,008
0,004
0
Adult brain
Mat > Pat
C.
p = 0.0775
0,006
0,0045
0,003
0,0015
0
ESC
D4
D6
D8
NPC
Bi.
D.
200 kb
mm9: chr15:72,587,105-72,891,273
Kcnk9
Trappc9
Chrac1
Ago2
Peg13
PE
35
H3K27ac
0
20
Maternal allelism
Paternal allelism
ChIP-seq
Fontal Cortex
20 kb
mm9: chr15:72,604,053-72,674,519
